# Classifier-guided Deep Oscillatory Neural Networks (cDONN) for capturing both Neural Dynamics and Behavior simultaneously

**DOI:** 10.1101/2025.06.27.661914

**Authors:** Sayan Ghosh, Sapna Raja, V. Srinivasa Chakravarthy

## Abstract

Generating EEG signals alongside behavioural actions introduces substantial biological complexity, akin to an abstract model that mimics rich oscillatory brain dynamics by simultaneously producing dynamic neural activity and corresponding behavioural responses. In this work, we propose a novel class of bio-inspired classifier guided Deep Oscillatory Neural Network (cDONN), designed to simultaneously model and capture both neural signal dynamics and behavioural patterns. The cDONN integrates a feedforward neural network with Hopf oscillator neurons, specialized in handling sequential data generation tasks. The model features a Y-shaped structure with two output branches: one for classifying the input image that signifies behaviour, and the other for generating corresponding EEG signals that represent neural dynamics. We evaluate the performance of the behaviour classification branch using the standard ramp classification accuracy used for the cDONN network. For the EEG signal generation branch, we benchmark the generated signals using classification accuracy via a pre-trained DONN signal classifier, alongside comparison with real EEG signals using metrics such as Inception Score, Frechet Inception Distance (FID), power spectrum analysis and topoplot analysis. Our analysis reveals a strong correspondence between the generated and real EEG data, particularly in the midline parietal region within the alpha frequency band, and in the left hemisphere within the delta band, indicating high fidelity and neurophysiological realism in the synthesized signals. Our approach offers a novel perspective on modelling the simultaneous relationship between neural activity and behaviour, pushing the boundaries of EEG signal generation and behavioural analysis.

## Introduction

In the last decade or two there has been a strong interest to develop large scale models of brain function. Ideally such models must be able to model both brain dynamics and its external manifestations in terms of behavior. However, there is an exclusive class of network models that can capture brain dynamics and another large class that can learn behavior. It remains an unresolved challenge to create models that can capture both brain dynamics and behavior.

There has been an extensive effort in leveraging networks of nonlinear oscillators to model large-scale brain activity. A prominent example is The Virtual Brain (TVB) platform, which utilizes large-scale oscillatory networks to simulate diverse functional brain dynamics observed in modalities such as EEG, functional Magnetic Resonance Imaging (fMRI), and Magnetoencephalography (MEG) [1]. For instance, Al-Hossenat et al. employed the Jansen-Rit (JR) neural mass model within the TVB framework to model slow-wave EEG activity (delta and theta bands) across eight brain regions, incorporating anatomical connectivity [2]. Similarly, another study used the same neural mass approach to reproduce alpha wave activity from four distinct brain regions [3]. In related work, a weighted mean field of a weakly coupled cluster of Hindmarsh-Rose (HR) neurons demonstrated near-synchronous dynamics, effectively reconstructing epileptic EEG time series [4]. Phuong and collaborators extended this line of research by employing networks of HR neurons and Kuramoto oscillators to model EEG signals in both healthy and epileptic states [5]. Moreover, similar nonlinear oscillator-based models, including Kuramoto and Hopf oscillators, have been successfully applied to simulate fMRI signals [6–7]. A variety of Hopf-based models have also been developed to explore and explain complex physiological phenomena, including cognitive functions, the sleep-wake cycle, schizophrenia, and Alzheimer’s disease [8–10].

Another modeling approach to brain dynamics involves use of deep learning models to generate EEG data. Several deep learning studies have explored reconstructing missing EEG channels using either the correlation with neighboring channels or deep learning approaches such as LSTM [11–12]. Among these, the method proposed by Bahador et al. [13], which leverages both temporal and spatial correlations, achieved the highest reported prediction accuracy of 82.48 ± 10.01%. EiDS is a novel emotion-inspired deep recurrent network that outperforms standard LSTMs in forecasting chaotic EEG time series by mimicking emotion-related brain structures for improved short- and long-term prediction [14]. hvEEGNet is a hierarchical variational autoencoder, built on EEGNet blocks and trained with a dynamic time warping loss, that delivers high-fidelity multi-channel EEG reconstruction [15]. To address the scarcity of high-quality EEG data for seizure prediction, a DCGAN is used to generate synthetic EEG spectrograms of preictal states [16]. EEG-GAN is a Wasserstein GAN-based framework with stability enhancements that generates realistic synthetic EEG time-series data for augmentation, super-sampling, and restoration of corrupted signals [17]. ATGAN is an attention-based temporal GAN designed to augment raw EEG time-series for personal identification—leveraging both temporal attention and GAN-based adversarial learning to boost identification accuracy by ∼7.8% on the BCI Competition IV-2a dataset [18]. Enhanced EEG Forecasting leverages a probabilistic WaveNet model to predict resting-state EEG in theta (4–7.5 Hz) and alpha (8– 13 Hz) bands up to ∼150 ms ahead—achieving mean absolute errors of ∼1 µV and outperforming autoregressive models by estimating both signal amplitude and phase while allowing uncertainty-guided filtering [19]. Also, current trends are to use diffusion models for EEG signal generation. Torma et al proposed Diffusion Probabilistic model (DPM) to generate synthetic EEG data [20]. In another study, authors introduced denoising diffusion probabilistic models (DDPMs) to generate synthetic EEG data [21]. Aristimunha, B et al. proposed latent diffusion models to create artificial sleep EEG data [22]. However, denoising diffusion probabilistic models (DDPMs) are useful to generate physiological time series like: electroencephalography (EEG), electrocorticography (ECoG), and local field potential (LFP)[23].

Standard deep learning models, particularly deep neural networks (DNNs), have demonstrated remarkable success in learning complex behavioral tasks such as vision, audition, and language processing. Convolutional neural networks (CNNs) have revolutionized visual recognition tasks by achieving human-level performance in image classification and object detection [24–25]. In the auditory domain, recurrent neural networks (RNNs) and more recently, transformer-based architectures have enabled accurate speech recognition and sound classification [26–29]. For language understanding, transformer models like Bidirectional Encoder Representations from Transformers (BERT) and Generative Pre-trained Transformers (GPT) have established new benchmarks in natural language processing [30–33], showcasing the ability of deep learning systems to generalize across diverse linguistic tasks. Multimodal models such as [34–37] have shown the ability to connect textual and visual understanding, performing tasks like image generation from text prompts and zero-shot image classification. Similarly, models like Whisper [38] have advanced speech recognition by being trained on multilingual, multitask datasets. These developments exemplify how increasingly sophisticated AI models are pushing the boundaries of machine perception and cognition, approximating complex human-like behavioral capabilities through end-to-end learning from large-scale data. These advances underscore the power of deep learning as a framework for modeling and replicating human-like cognitive and perceptual abilities. Although at the level of input/output the above class of networks showed excellent performance, the internal neural dynamics of these networks are rather weak in neurobiological plausibility. In real brains, collective neural activity exhibits rich oscillatory dynamics which is expressed in terms of frequency bands like alpha, beta, gamma etc [39]. Deep neural networks, of both feedforward and recurrent kind, do not exhibit such oscillatory dynamics. Thus, the two broad classes of models described above effectively capture large scale brain dynamics, both in normal and disease conditions, their inherent limitation lies in their inability to learn behavior. There is a rapidly growing interest in developing large-scale models of brain function that can serve as powerful tools for understanding the complex interplay between neural dynamics and behavior. Ideally, these models should not only replicate the rich, nonlinear dynamics of neural activity across anatomically and functionally distinct brain regions, but also faithfully reproduce the brain’s external manifestations ranging from perception, cognition, and emotion to goal-directed behavior and motor actions. Such integrative models hold the potential to unravel the mechanisms underlying brain function and dysfunction, offering profound insights into both normal and pathological states, and ultimately enabling more precise and personalized approaches in neuroscience and neuroengineering.

With this motivation, we propose a novel, biologically plausible Classifier guided Deep Oscillatory Neural Network (cDONN) for a generative AI task. Our network integrates two core branches: one that classifies stimulus images into 10 distinct categories and another that generates corresponding EEG signals from the same images. This hybrid architecture leverages multiple DONN modules to simultaneously perform classification (behavior) and signal generation (dynamics). To the best of our knowledge, no existing work has addressed this type of integrated model that mirrors the brain’s ability to handle both behavioral responses and dynamic neural activity.

The outline of the paper is as follows. Section 2 presents detailed Methods, specifications, and the data sets used in section 2.1, The network architecture is explained in section 2.2. Section 3 describes the results which are discussed in the final section. At last, we have a discussion, conclusion and future direction section.

## 2. Methods

### 2.1 Dataset

The present study utilized a dataset comprising three categories: Digits, Characters, and Objects [40]. EEG data were recorded using 14 channels (AF3, F7, F3, FC5, T7, P7, O1, O2, P8, T8, FC6, F4, F8, AF4) from the Emotiv EPOC device, following the 10-20 electrode placement system. The dataset was collected from 23 participants at a sampling rate of 2048 Hz, which was later down-sampled to 128 Hz. The dataset includes EEG recordings corresponding to three visual stimuli: digits (0-9), English characters (ten classes), and objects selected from the ImageNet dataset [41]. Each visual stimulus was displayed for 10 seconds, followed by a 20-second relaxation period. Following the preprocessing steps outlined in [42], the dataset was segmented using a 0.25-second sliding window with an overlap of 0.0625 seconds. Additionally, z-score normalization [43] was applied to standardize the EEG data.

### 2.2 Classifier-Guided networks

This classifier-guided network (fig. 1 and fig. 2(A-C)) comprises two main modules: (A) the Image Classifier network and (B) the Signal Generation network. Figure 1 presents an abstract overview of the entire study, while Figures 2(A–C) provide a more detailed depiction of each module within the classifier-guided network. In Fig. 1, we can see a fork-shaped model where one side represents the behavioral output (Input to Image Classifier) and the other side represents the signal generation part. Whereas a more detailed figure (2(a)) shows the behaviour part (image classification), and Fig. 2(B)-2(C) describes the dynamics (signal generation) part.

**Figure 1:**
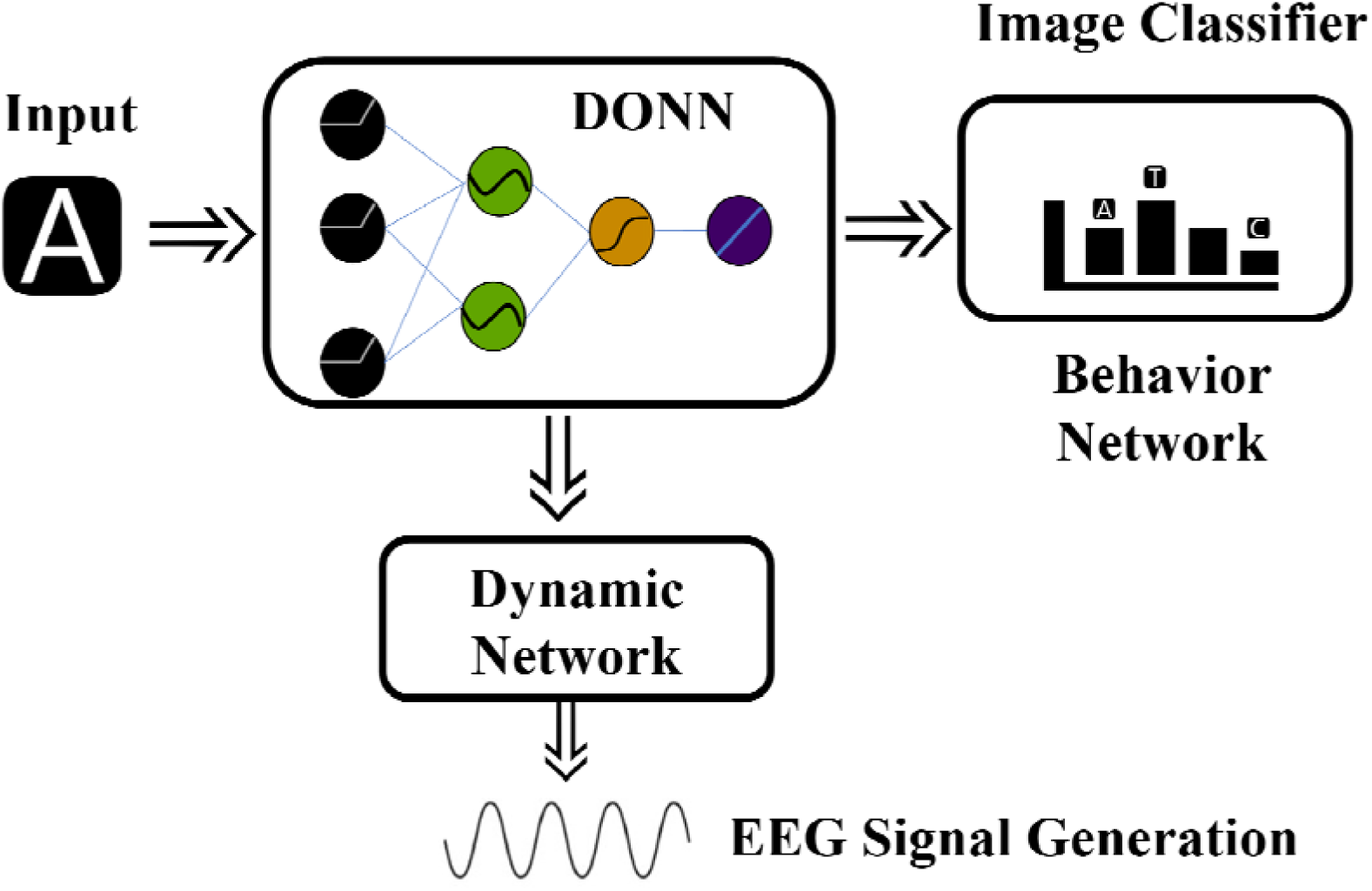
Abstract flow diagram of Classifier-Guided Network.

**Figure 2(A-C):**
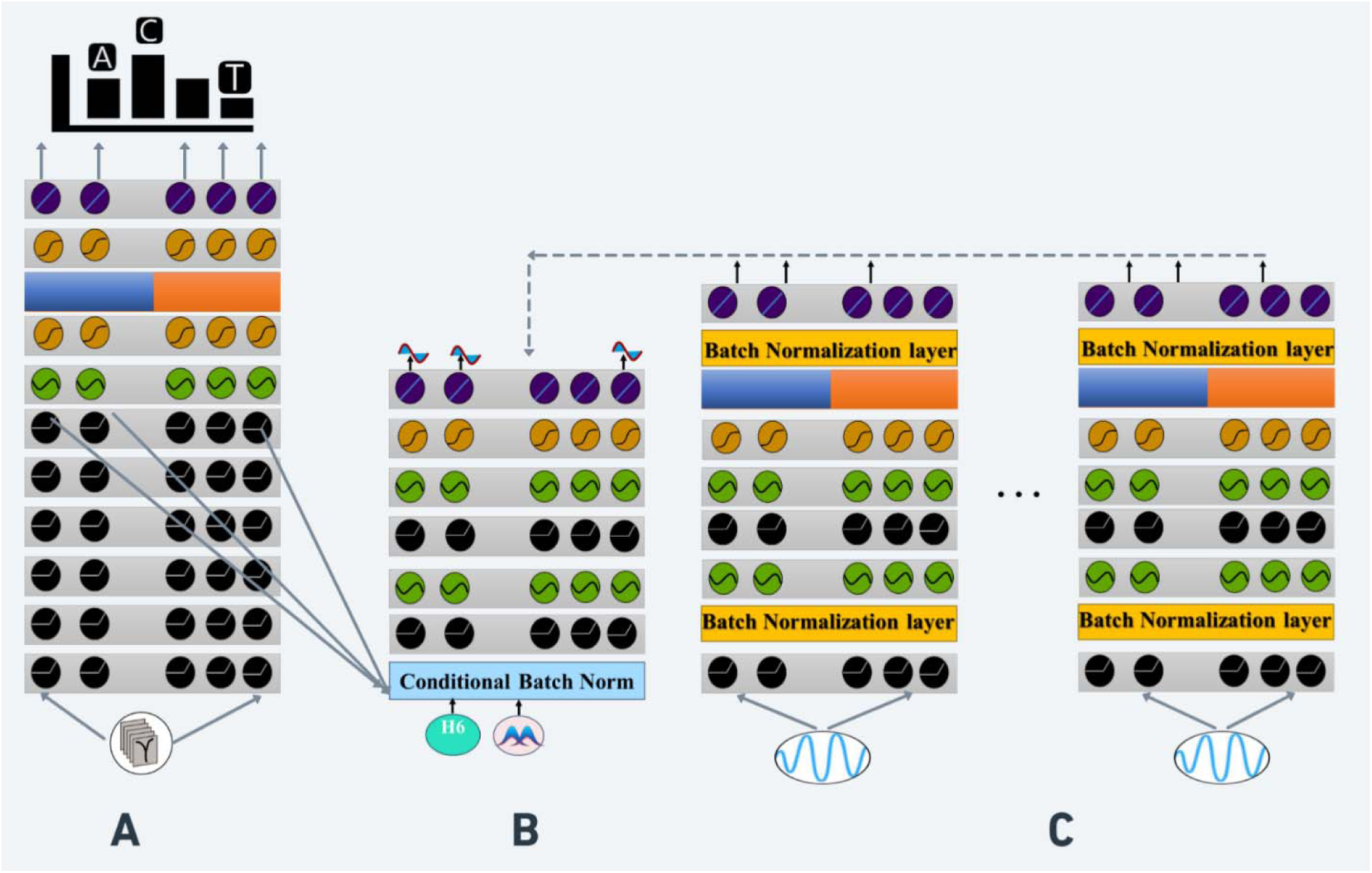
Detailed architecture of the classifier-guided Deep Oscillatory Neural Network. (A): Image classifier Network, (B): Signal Generator Network, (C): Signal evaluator Network

Additionally, the Signal Generation network includes a sub-branch (fig 2C) called the Signal evaluator network, which aids in learning the EEG signals in network (B) associated with specific stimuli. All three networks (A, B, and C) are fundamentally built upon the Deep Oscillatory Neural Network (DONN), a modeling framework developed by our group [42, 44–45]. Each DONN is composed of a series of Hopf oscillatory dynamic neuron layers, followed by nonlinear static neuron layers such as ReLU, sigmoid, and tanh [42, 44–45]. In this joint network, the Signal Evaluator network and Signal Generation network are trained jointly, with the central component being an integrated architecture that includes an Image Classifier and a Signal Generation network for synthesizing EEG signals based on visual input. The visual stimuli classifier and the generator share common initial layers. From the next paragraph, we will describe each of the networks.

### 2.3 Image Classifier Network

The architecture of the image classification network is illustrated in Figure 3. This general feedforward network design is suitable for a wide range of classification tasks [42, 44–45]. The network begins with a fully connected (all-to-all) stage from the input layer to the first hidden layer (H1), which consists of Complex ReLU (cReLU) neurons. All neurons and their corresponding weights in the network are complex-valued. Then there are six consecutive hidden layers (H1 to H6), all using the cReLU neurons.

**Figure 3:**
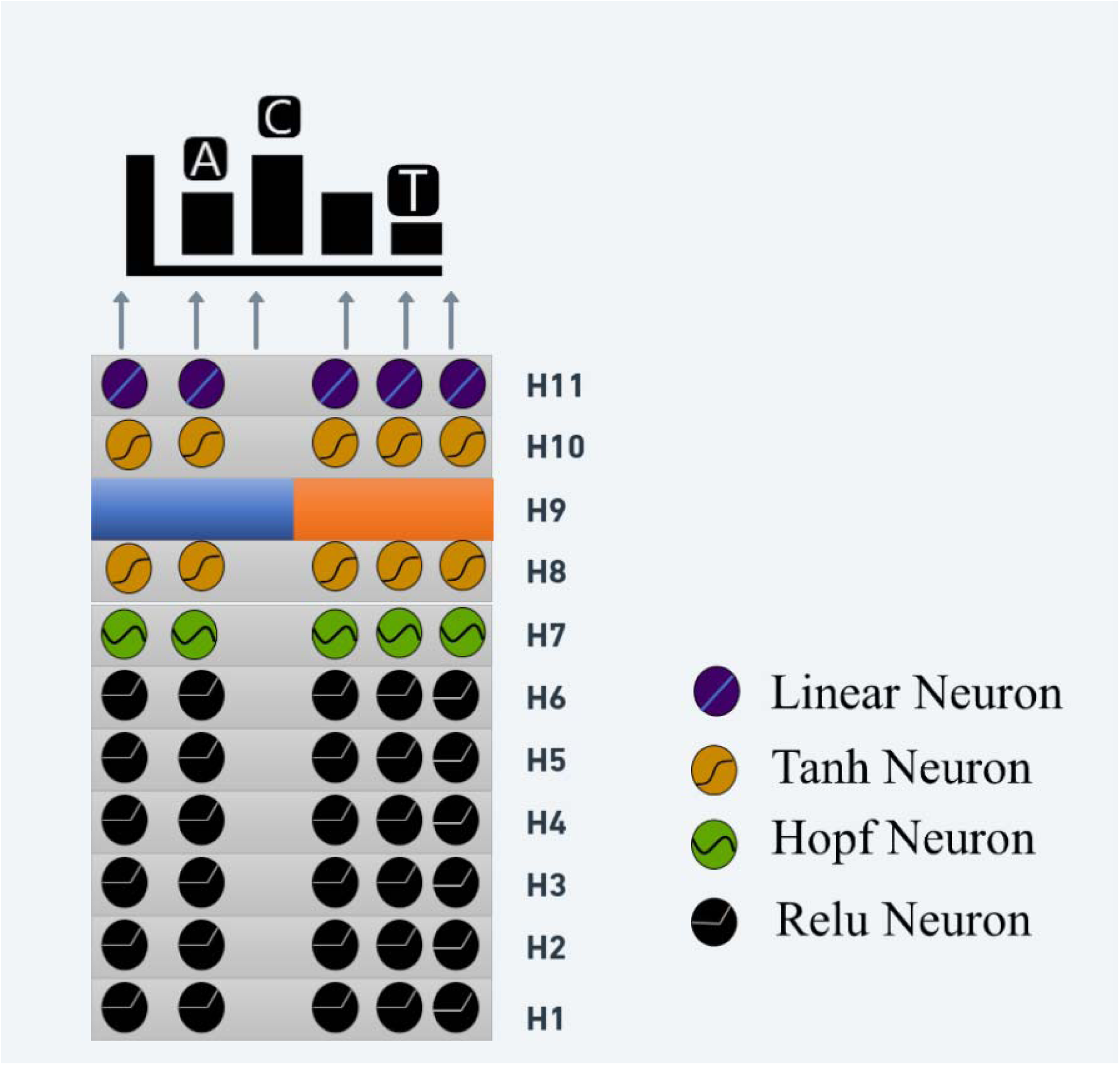
Image classifier Network.

A complex-valued weight matrix connects the image input to the first hidden layer H1. The resulting product is then processed through the complex ReLU (cReLU) activation function, as defined in Equations 1.2–1.4. This activation function applies the standard Rectified Linear Unit (Relu) separately to both the real and imaginary components of. The complex weight matrix can be expressed as:

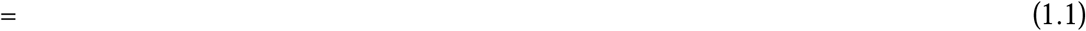

where 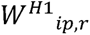 is the real part of connecting weight between the input to H1 layer, 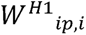 is the imaginary part of connecting weight between the input to H1 layer. The complex feed forward weight of H1 layer has been shown in (1.1).

Relu is applied to the 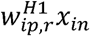 and 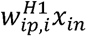

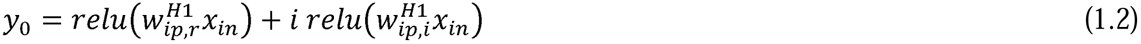

where normal relu() can be defined as:

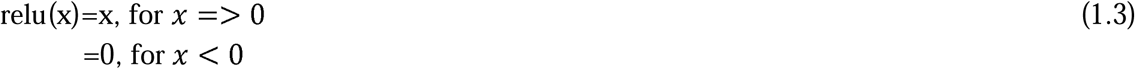

Equation (2.1) can be written as:

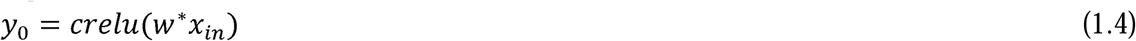

which defines a complex-valued ReLU or a crelu(). *X_in_* is the input to the H1 layer. The forward pass computations from H1 layer to H6 layer can be described as above equn. 1.4. Then the output of the H6 layer (cReLU layer) is presented as input to the H7, which is a layer of Hopf oscillators.

The oscillatory neuron implemented in the H7 layer is based on the Hopf oscillator [44], a type of harmonic oscillator characterized by a stable limit cycle. The dynamics of the standard Hopf oscillator are governed by the following complex-valued differential equation,

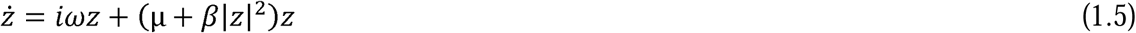

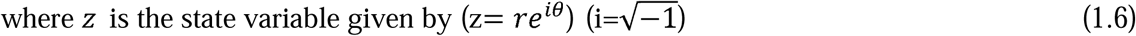

The polar coordinate representation of the above oscillator equation is:

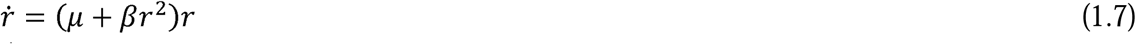

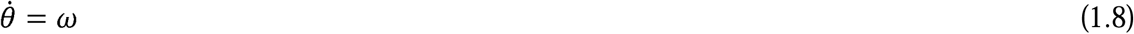

Here, *r* and (*θ* represent the state variables of the oscillator. The term (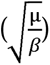 defines the amplitude of oscillation, where μ & β are the bifurcation parameters that control the system’s behavior. The parameter ω denotes the natural frequency of the oscillator. The external input signal is given by I(t), and ω_0_ represents the angular frequency of the externally applied stimulus.

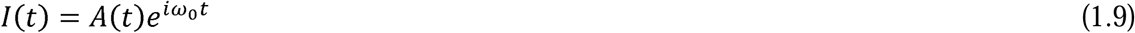

where the input presented as an additive input to the oscillators is shown below:

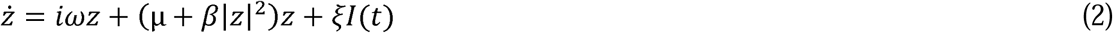

The polar coordinate representation of the last equation is:

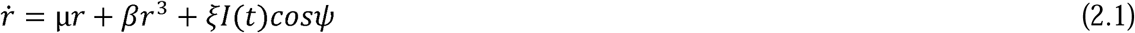

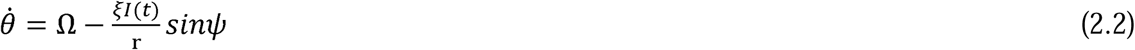

where 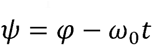 and 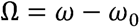 is the difference between the angular frequencies of the the oscillators and the external input.

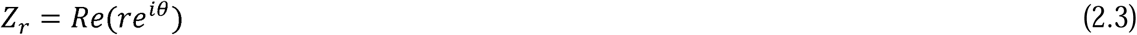

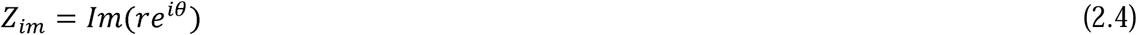

Here, Z_r_ and Z_im_ denote the real and imaginary components of the oscillator’s state variable in In this formulation, μ & β are the bifurcation parameters, ω is the natural frequency, and the complex domain. The variable z represents the complex-valued state of the Hopf oscillator. I(t)refers to the external input signal applied to the oscillator. In our previous study we have described the network of hopf oscillatory neuros [44].

Note that the connection between H6 ReLU layer and H7 oscillatory layer is one-to-one, whereas all other previous weight stages are all-to-all connections. Then we use a Complex Tanh (cTanh) neuron layer(H8). The frequencies _j_ of the oscillators in the H7 oscillator layer are randomly initialized within the typical EEG frequency range of 0.1 to 64 Hz. In contrast to the approach described in [46], the oscillators in this layer operate independently, with no lateral connections between them.

All the feedforward weights and the oscillator’s frequency are trained using TensorFlow Autograd. At the H9 layer we have concatenated the real and imaginary parts of the weight. Subsequently, we have two more layers, H10 and H11, having Tanh and linear activation functions, respectively. Graphical illustration of Image classifier network has been shown in fig 3, which is a subpart of Fig. 2(A-C).

The Image Classifier network is designed for classification tasks, with the number of output neurons implemented as linear neurons matching the total number of classes. All hidden layers consist of complex-valued neurons, except for the H11 layer, which contains real-valued neurons. Signal classification is achieved by defining the target output within a supervised learning framework, following the approach outlined in our previous DONN model [44].

For vector classification, since the task can be completed in a single step, the desired output is represented using a standard one-hot encoding. In contrast, signal classification requires analyzing the input over a finite duration, making one-hot encoding unsuitable for the entire duration. Instead, the target output for the neuron corresponding to the correct class is defined as a ramp signal, while the outputs of the remaining neurons remain at zero throughout the time (see eqns. 2.6-2.7).

For visual input image classification, we have used Ramp MSE loss (eqn 2.5).

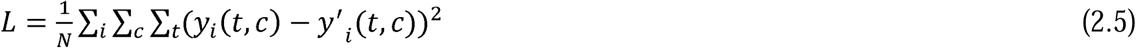

Where *y_i_* (*t, c*) and *y’_i_* (*t, c*) represent the true ramp signal and predicted signals for the c^th^ class at time point t for sample i. The loss is computed as the mean squared error over all samples, classes, and time steps. Specifically, the loss is averaged over *N*, where *N* is the total number of terms in the sum, given by the product of the number of samples in a batch, the number of time points, and the number of classes:

N=(number of samples)×(number of time steps)×(number of classes)

This formulation ensures that the loss equally accounts for temporal dynamics, class-wise predictions, and variation across the batch.

The target output for the neuron corresponding to the correct class is defined as a ramp signal, Thus the target ramp *y_i_* (*t, c*) can be defined (eqns 2.6-2.7): while the outputs of the remaining neurons remain at zero throughout the presentation.

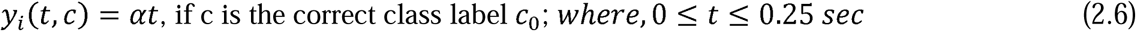

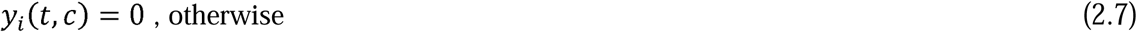

*y_i_* (*t, c*) is a (32×10) matrix where each row corresponds to a time point t and each column to a class ccc. For a given sample i, only one column—corresponding to the target class k—contains a non-zero signal. The entries in the k-th column increase linearly from 0 to 0.25 across the 32 time points, forming a ramp signal. All other columns are filled with zeros. The slope α of the ramp is set to 8, and is used to generate the target ramp profile:

### 2.4 Signal Generator Network

To introduce variability in the generated EEG signals, we inject noise into the Signal Generator Network. This noise is modeled using a multi-dimensional Gaussian Mixture Model (GMM) [47], consisting of three component Gaussian distributions (Z1, Z2, Z3), as described in Equations (2.1–2.5). These Gaussians are summed to form a richer and more robust noise representation.

To condition the signal generation process, we use a Conditional Batch Normalization (CBN) mechanism. Specifically, the hidden layer output H6 of the Image Classifier Network (Fig. 3)— referred to as Network A—is used to modulate the injected noise via CBN (Fig. 4). This allows the generator to adaptively shape the EEG signal based on the visual features captured in H6, enabling conditional signal generation. The CBN-based modulation introduces meaningful variability in the output, helping the model better capture the diversity of EEG patterns corresponding to different images.

**Figure 4:**
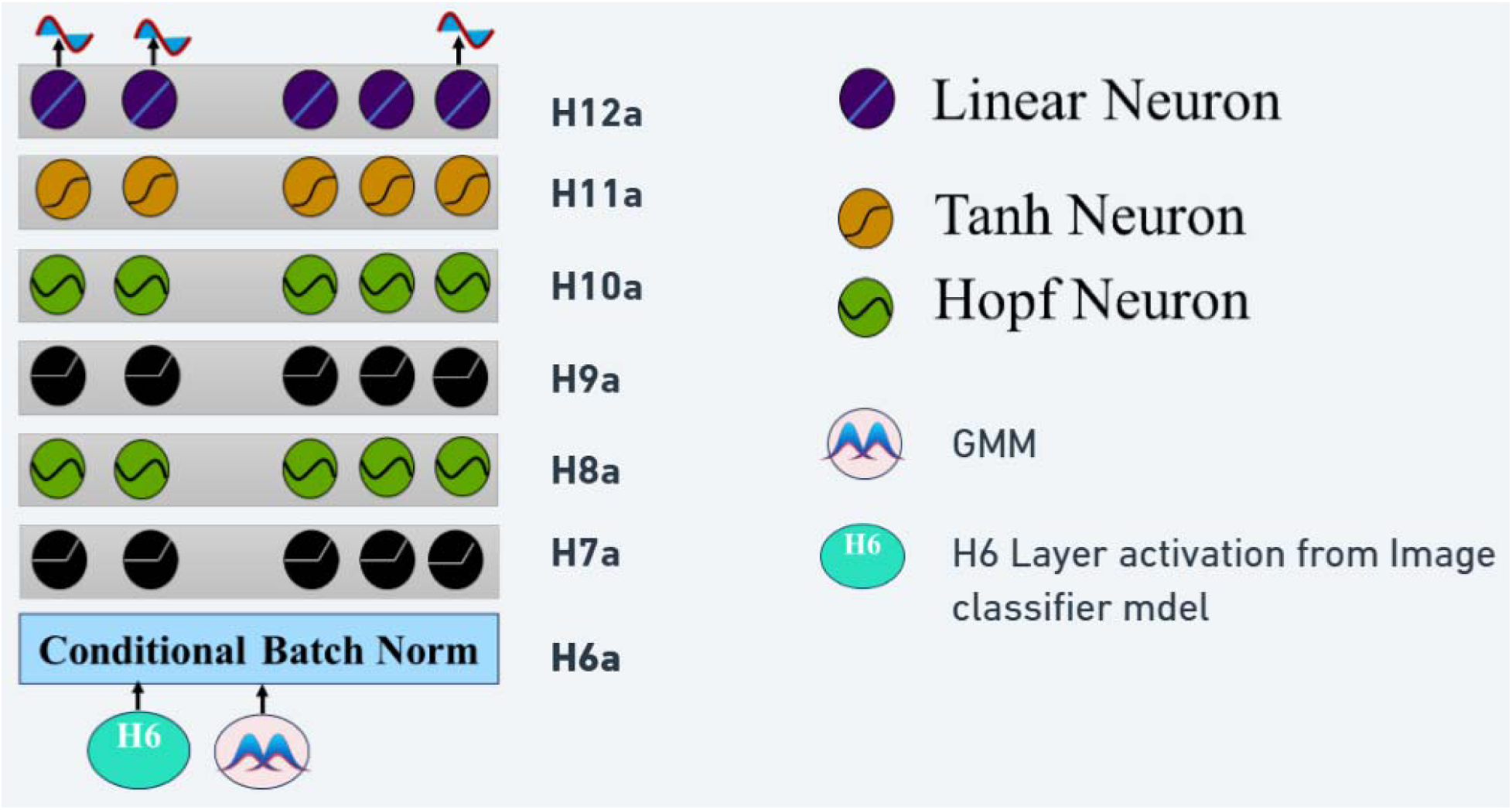
Signal Generator Network

In order to produce EEG signal variability, we introduce noise in the Signal Generator Network. The noise is modeled as Gaussian Mixture [59] consisting of three component gaussians as described in eqns (2.1-2.5). We used conditional batch normalization approach to introduce the noise into the model, which eventually achieved the required variability in the generated data, at the output layer (H12a). The hidden layer output (H6) of the Image classifier network (fig. 3) is sent to the generator network in addition to the Gaussian Mixture noise (fig. 4). In this work we used H6 layer output of network A. The noise is tuned by the H6 layer output of image classifier network. Graphical illustration of Signal generator network has been shown in fig 4, which is a subpart of Fig. 2(A-C)

Where,

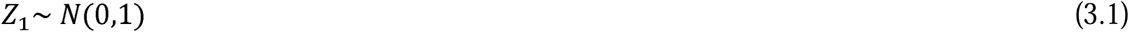

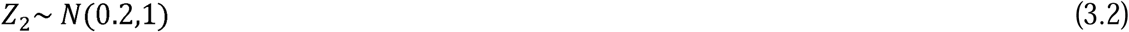

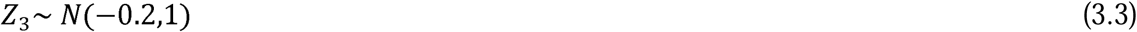

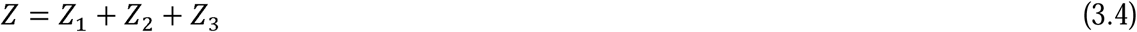

Let *h*_6,*r*_ *h*_6*i*_ are the H6 layer output. After applying conditional batch normalization with the gaussian mixture noise (Z):

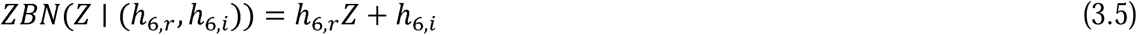

After the Conditional Batch Normalization(H6a), the output (ZBN) is passed to the H7a layer, which consists of cReLU neurons. The activation from this layer is then forwarded to the H8a layer, composed of Hopf oscillatory neurons. This cReLU-Hopf pair operation is repeated once more through layers H9a and H10a. All cReLU-Hopf pairs use one-to-one connections, while all other layer connections are fully connected (all-to-all).

Following the H10a Hopf layer, the signal output is concatenated and passes through two consecutive layers: cTanh layer (H11a), and finally, a linear layer (H12a). In this generator network, the target output is the raw EEG signal.

The loss function of the Generator Network can be written by (eqn. 3.6)

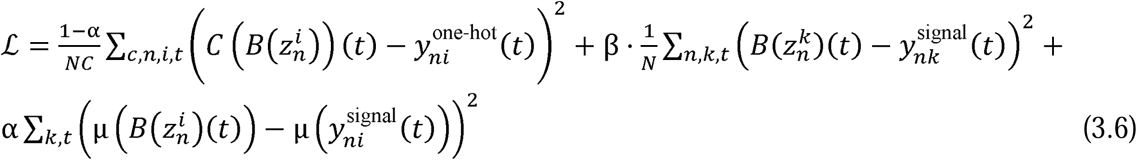

where, *α,β* are the weighting parameters. *α* = 0.99, *β* = 0.01.

C number of selected EEG evaluators; N is the number of signals in a batch

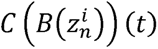 ; where 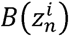 is the EEG signals generated by the EEG signal generator, 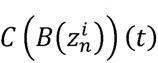 is the output of the randomly selected subset from the bank of evaluators on the generated signal

The target output 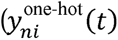 from each classifier can be defined (eqns 3.7-3.8):

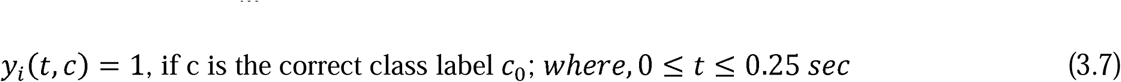

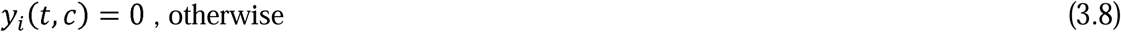

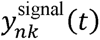 is the true EEG signal.

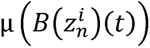 is the mean of the generated signals from a batch,

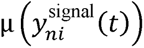 is the mean of true EEG signal.

In eqn. (3.7), the 1^st^ term denotes Classifier Loss from selected classifiers, the 2^nd^ term denotes MSE loss between true and generated signals, the 3^rd^ term distribution matching loss function aiming to minimize the batch mean between true and generated signals.

### 2.5 * Bank of signal evaluator networks

The weak classifier signal evaluator network consists of seven individual signal evaluator modules, each based on a simplified DONN architecture. These modules take either real or generated EEG signals as input and perform a ten-class signal classification task [42]. Compared to the standard DONN classifier, these evaluators have significantly fewer parameters and neurons in each layer to prevent overfitting and to ensure that the ensemble of networks learn to recognize distinct characteristics of the EEG signals such as spectral or temporal patterns.

The design of these weak evaluators ensures diversity in learning, so that each module contributes uniquely to guiding the signal generation process. Collectively, these evaluators form a robust system that provides gradient feedback to the signal generator network, helping it learn to produce realistic and class-representative EEG signals.

In each training epoch:

- Each evaluator is also trained on randomly selected batches of real EEG data, encouraging them to generalize across different data distributions.
- The signal generator network produces synthetic EEG signals, which are passed to a subset of ramdomly selected 5 out of seven evaluator networks, which in turn provide feedback to the generator network.

The weak signal evaluator network comprises a set of 7 individual signal evaluator modules, each of which comprises a similar DONN architecture, and either takes in the real/generated EEG signals as input and performs a ten-class signal classification task [42] These networks are designed in such a way that number of neurons in each layer is much lesser than number of neurons in strong classifier modules. This to to ensure that each evaluator doesn’t overfit on the training data and learn to classify EEG signals based on different patterns (Spectral/Temporal characteristics). A system of such weak evaluator networks collectively acts like a strong driving agent guiding the signal generator network to generate representative EEG signals for each class. A single signal evaluator module is illustrated fig. 5. The output of each evaluator is a one hot signal. Five of these seven networks are randomly selected to drive the signal generator network in each epoch of training. In each epoch of training, the individual evaluator networks are trained using randomly selected subset of real EEG data, so that collectively the system of weak signal evaluator networks guides the signal generator network to generate signals sampled from different modes of the data.

**Figure 5:**
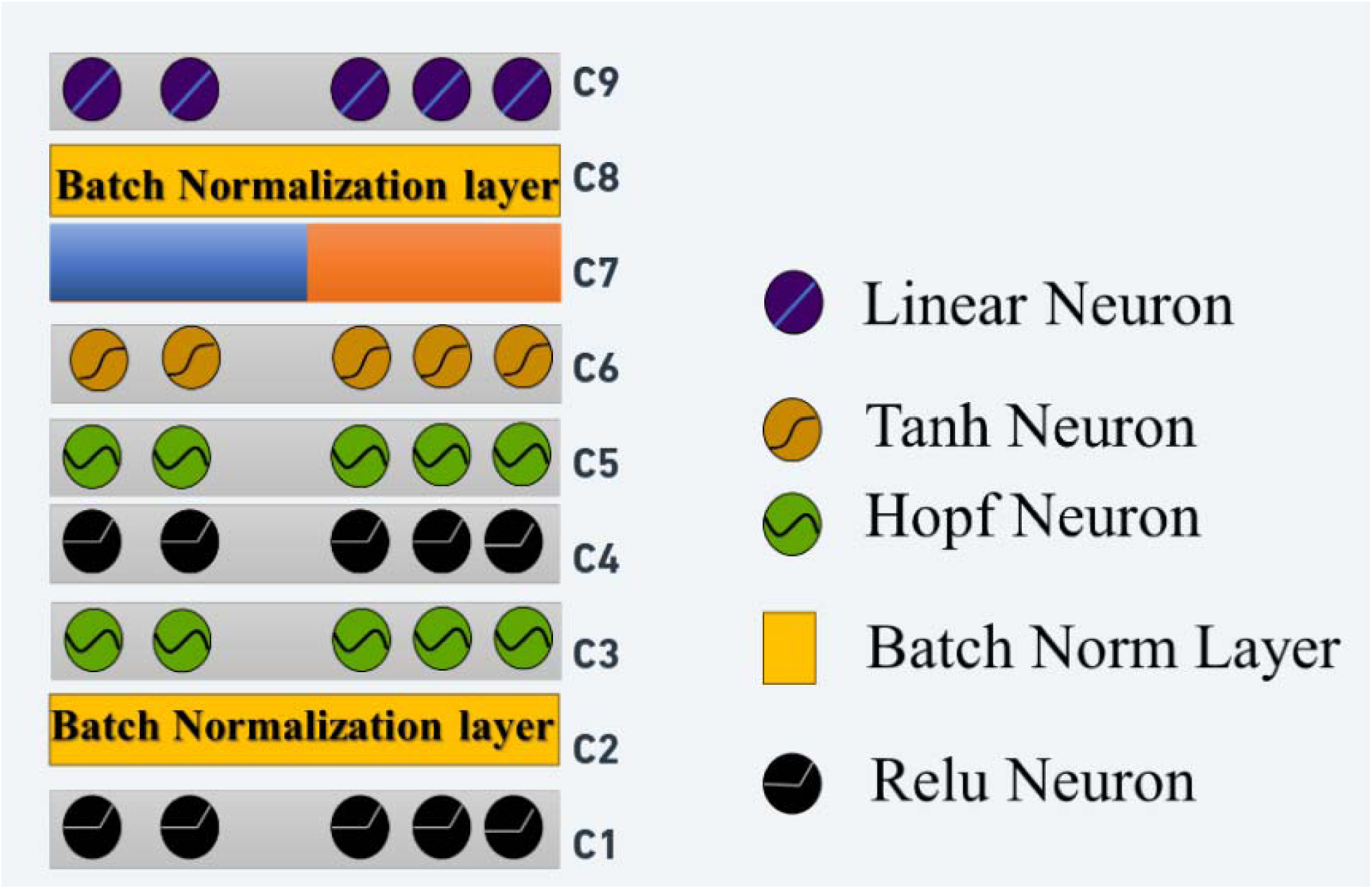
Signal evaluator Network, which is a subsection fig (2(C))

In this architecture, the EEG signals are first passed through a cReLU layer (C1), followed by batch normalization (C2). Next, the output is processed by a Hopf oscillatory neuron layer (C3). This is followed by another pair consisting of a cReLU layer and an oscillatory neuron layer (C4–C5). After that, a cTanh activation layer (C6) is applied. The real and imaginary components of the C6 layer’s output are then concatenated. This concatenated output undergoes another round of batch normalization, and finally, it is passed through a linear layer (C9) to produce the output.

### 2.6 Training loop

The training process begins with the K signal classifiers, where each classifier is presented with EEG signals from distinct batches. This ensures that the classifiers learn different patterns or subspaces spanned by signals of a specific class, thereby preventing mode collapse. After the classifiers are trained, the generator is subsequently trained. During its training, the generator receives feedback from any randomly selected m out of the K classifiers. Additionally, it incorporates other loss components, including distribution matching and mean squared error (MSE) loss.

#### Algorithm: Training of Generator and EEG Signal Evaluators

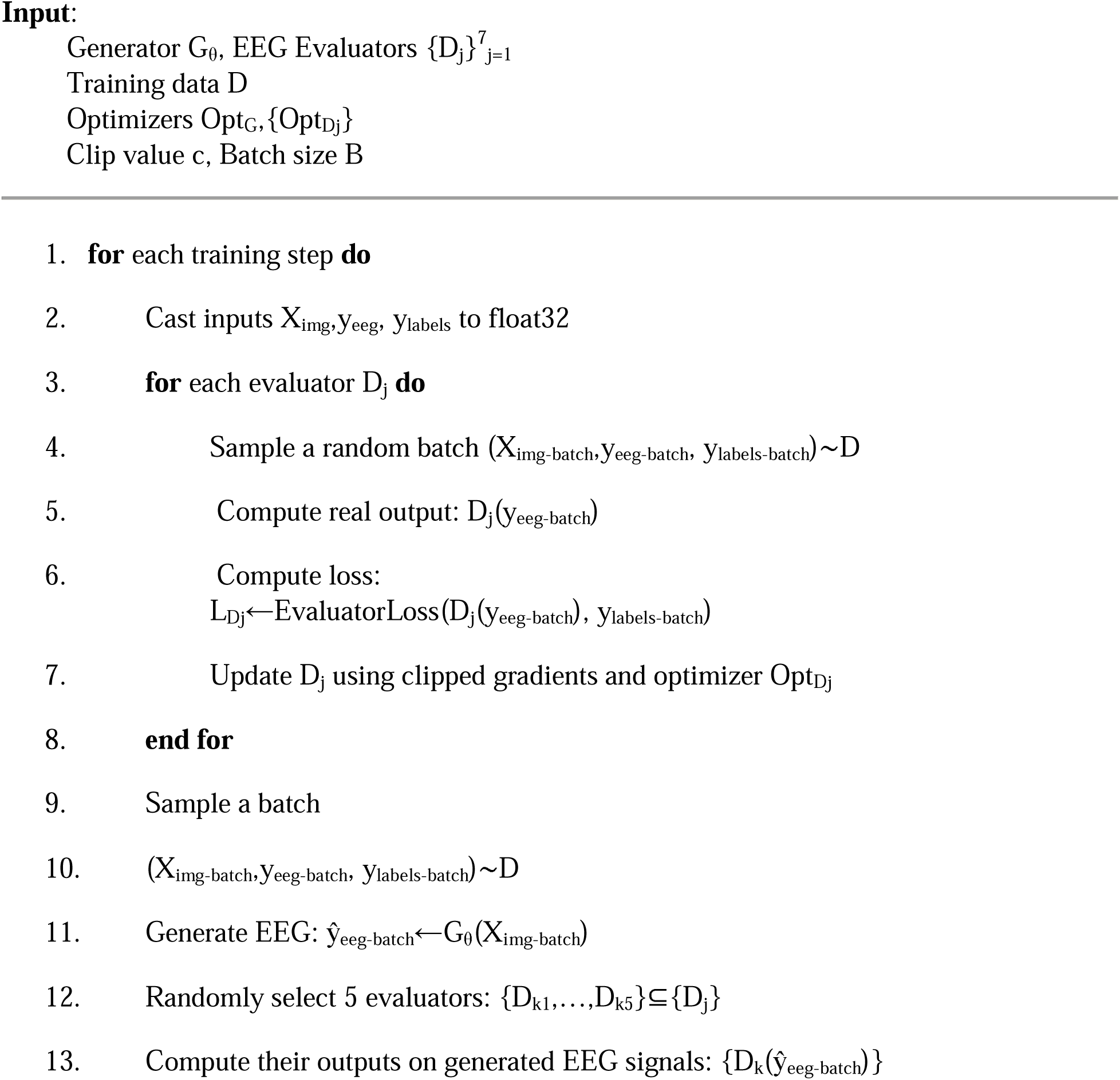
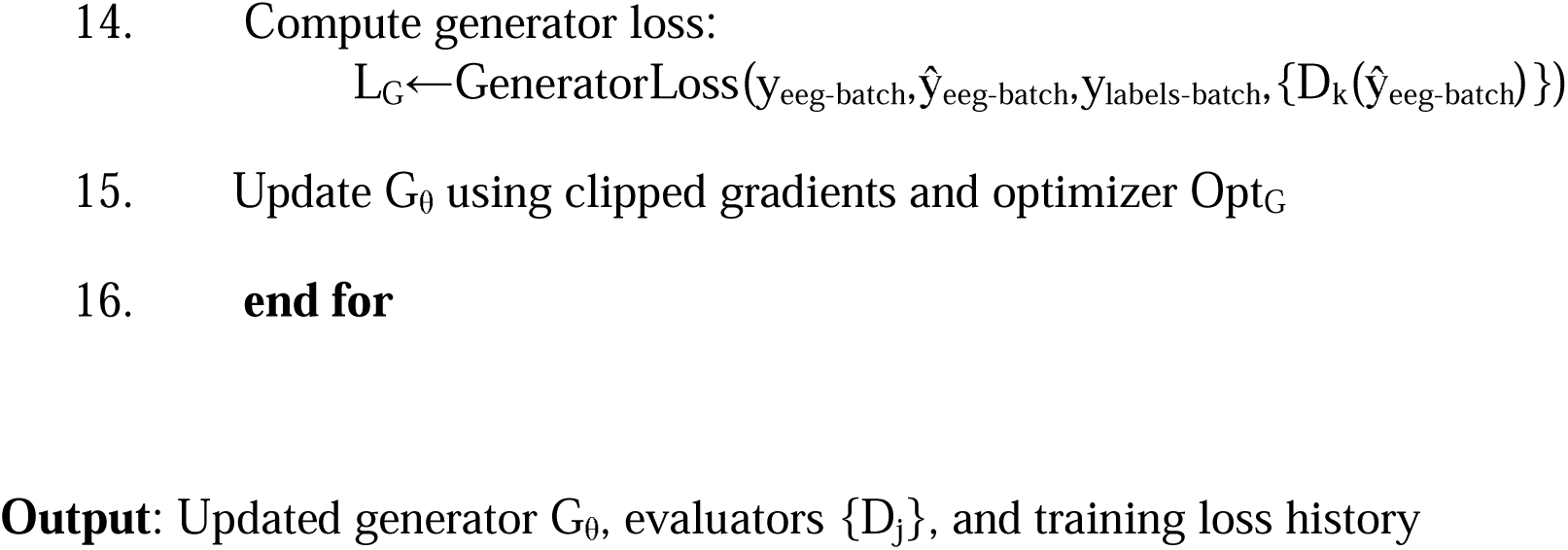

Where *G_θ_* is EEG generator; *D_j_* is one unit of EEG Evaluator,

Training Data {*D*} represents all sorts of training data used in the model, comprising of character images, EEG signals and the desired labels.

*Opt_G,_ Opt_D_* are the optimizers for EEG signal generator and EEG evaluator. Batch size B is used for training.

c is the clip to the gradient so that it does not exceed 1.

{*D*} =*X_img_*: stimulus images, *y_eeg_*: corresponding EEG, *y_labels_*: desired ramp output.

Select a random batch of images (*X_img,batch_*), EEG signal (*y_eeg,batch_*) and the corresponding desired output (*y_labels,batch_*) for the signal evaluator module. The output for an individual signal evaluator module is: *D_j_*(*y_eeg,batch_*). 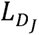 is the evaluator loss which is calculated based on *y_eeg,batch_* and *y_labels,batch_*.

For EEG generator (*G_θ_*) network: *ŷ_eeg,batch_* is *G_θ_* produced signal. *L_G_* is the EEG generator loss function.

{*D*l,*D*2,*D*3,…*D*7} are the seven signal evaluator modules.

### 2.7 Pre-trained DONN Classifier

Earlier we demonstrated the classification of ThoughtViz EEG data using the DONN model [42, 44]. In the present study, a pre-trained DONN network (Fig. 6), was employed for EEG classification [42], with its architectural specifications and parameter settings summarized in Table (1). The signals generated by the Signal Generator Network were evaluated using this pre-trained DONN model, enabling a consistent and reliable assessment of the generated signal quality within the same classification framework.

**Figure 6:**
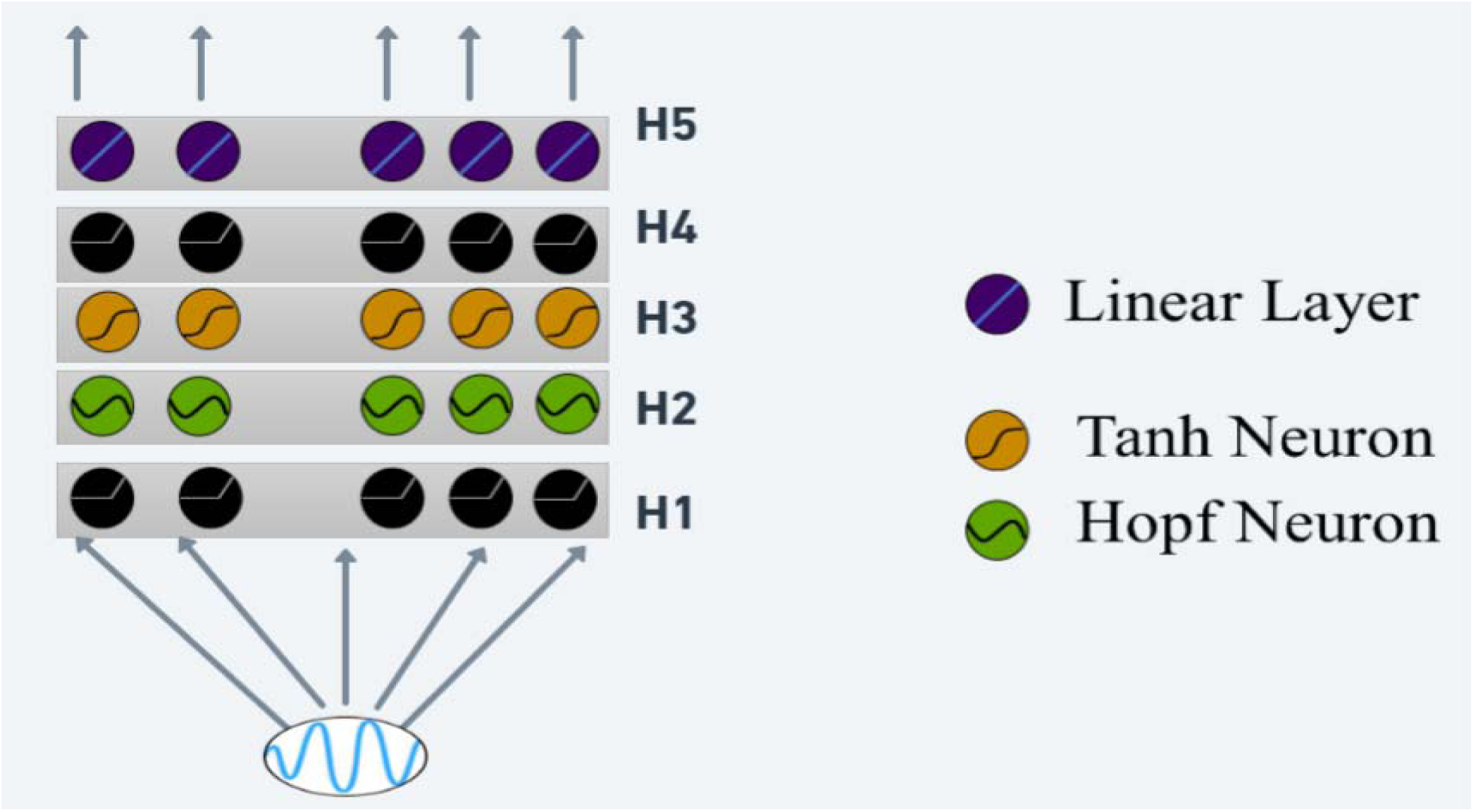
Pre-trained DONN Model

## 3. Results

### 3.1 Image Classifier Network

In the ThoughtViz dataset, each character is represented by 800 distinct font variations. For our analysis, we used 4000 EEG samples per character. To ensure alignment between the EEG signals and the corresponding stimulus images, we repeated each of the 800 font variants five times, resulting in a balanced set of 4000 image samples per class. However, the dataset does not provide information about which specific font was used in each sample of EEG.

For the testing phase, we used a set of 540 samples of EEG for each class, where each character includes 100 font variations, each font repeated 54 times, producing 540 image samples per class. This dataset was used in the testing phase (Table 2).

**Table 1.** Parameters of the pre-trained DONN architecture [47].

| Model parameters | Details |
| --- | --- |
| Input signal size | 32*14, where 32 is the number of time points, 14- channel |
| H1 Layer | ReLU neuron, units-250 |
| H2 | Hopf oscillators, units-250 |
| H3 | Tanh neuron, units-128 |
| H4 | Tanh neuron, units-64 |
| H5 | linear, units-10(10 class problem) |
| Hopf oscillator frequencies | [0.1-64 Hz] |
| Learning rate | 0.01 |
| Optimizer | adam |
| Number of epochs | 250 |
| Oscillator external input scaling factor ( ) | 4 |

**Table 2.** Training, testing, and splitting of the Image and EEG.

| Dataset | Train/ Test Image sequences | Train/ Test EEG |
| --- | --- | --- |
| Training (Individual class) | 4000*32*20*20*1 | 4000*32*14 |
| Testing (Individual Class) | 540*32*20*20*1 | 540*32*14 |

The original images are of size 20×20 pixels, which we flattened into 400-dimensional vectors before presenting them to the classifier network. Since our model is oscillatory and the EEG data consists of 32 time points, we stacked each image vector across all 32 time points to match the temporal structure of the EEG signals. Table 2 provides a detailed summary of the training and testing samples of both images and EEG signals used for each character. Since our image classification network is also based on the DONN architecture (Fig. 3), the ‘H7’ layer is implemented using a Hopf oscillatory network. The oscillatory parameters for the H7 layer are detailed in Table 3. Specifically, the natural frequencies (ɷ) of the Hopf oscillators are initialized within the ThoughtViz EEG signal frequency range of 0.1–64 Hz.

**Table 3.** Oscillator parameters used in all 3 networks.

| Oscillator Parameters | Signal Generator Network | Image classifier Network (H7 layer) | Classifier Networks |
| --- | --- | --- | --- |
| Oscillator external input scaling factor ($\epsilon$) | 100 | 2 | 2 |
| Hopf oscillator frequencies ($\omega$) | 0.1-64Hz | 0.1-64Hz | 0.1-64Hz |
| Range of omegas at $t=0$ | 0-pi/4 | 0 | 0 |
| Range of r at $t=0$ | 0-0.5 | 1 | 1 |
| Bifurcation Parameter ($\beta$) | 1 | 1 | 1 |
| dt (sec) | 1/128 | 1/128 | 1/128 |

The image classifier network is trained separately first with a standard DONN. The image classifier processes a 20×20 image, which is flattened into a 400-dimensional vector and stacked 32 times along the temporal axis, resulting in an input of shape (batch size, 32, 400). The image classifier network then incorporates an oscillatory layer with 25 neurons, followed by a ReLU layer with 15 units, and concludes with a final linear layer producing an output of 10 units for classification (fig 3). Finally, a ramp based desired target is used at H11 layer of image classifier network (fig 3). Model parameters for the Image classifier network are shown in Table 4.

**Table 4.** Image classifier network Parameters.

| Model Parameters | Values |
| --- | --- |
| H1 | 250 ReLU Neurons (Common Layer) |
| H2 | 200 ReLU Neurons (Common Layer) |
| H3 | 150 ReLU Neurons (Common Layer) |
| H4 | 100 ReLU Neurons (Common Layer) |
| H5 | 50 ReLU Neurons (Common Layer) |
| H6 | 25 ReLU Neurons (Common Layer) |
| H7 | 25 Hopf Neurons (Classifier) |
| H8 | 20 Complex Tanh Layer (Classifier) |
| H9 | 15 Complex Tanh Layer (Classifier) |
| H10 | 10 Linear Neurons (Classifier) |

### 3.2 Signal Generation network

The signal generator network builds upon the three shared layers from the Image classifier network, followed by two pairs of cRelu-Hopf oscillator layers, a tanh activation layer and a linear activation layer. The shared portion of the network consists of six ReLU layers, which reduce the dimensionality of the 400-dimensional input image to a compact 25-dimensional vector, effectively encoding the latent information from the image. The detailed model parameters for signal generator network is shown in Table 5.

**Table 5.**
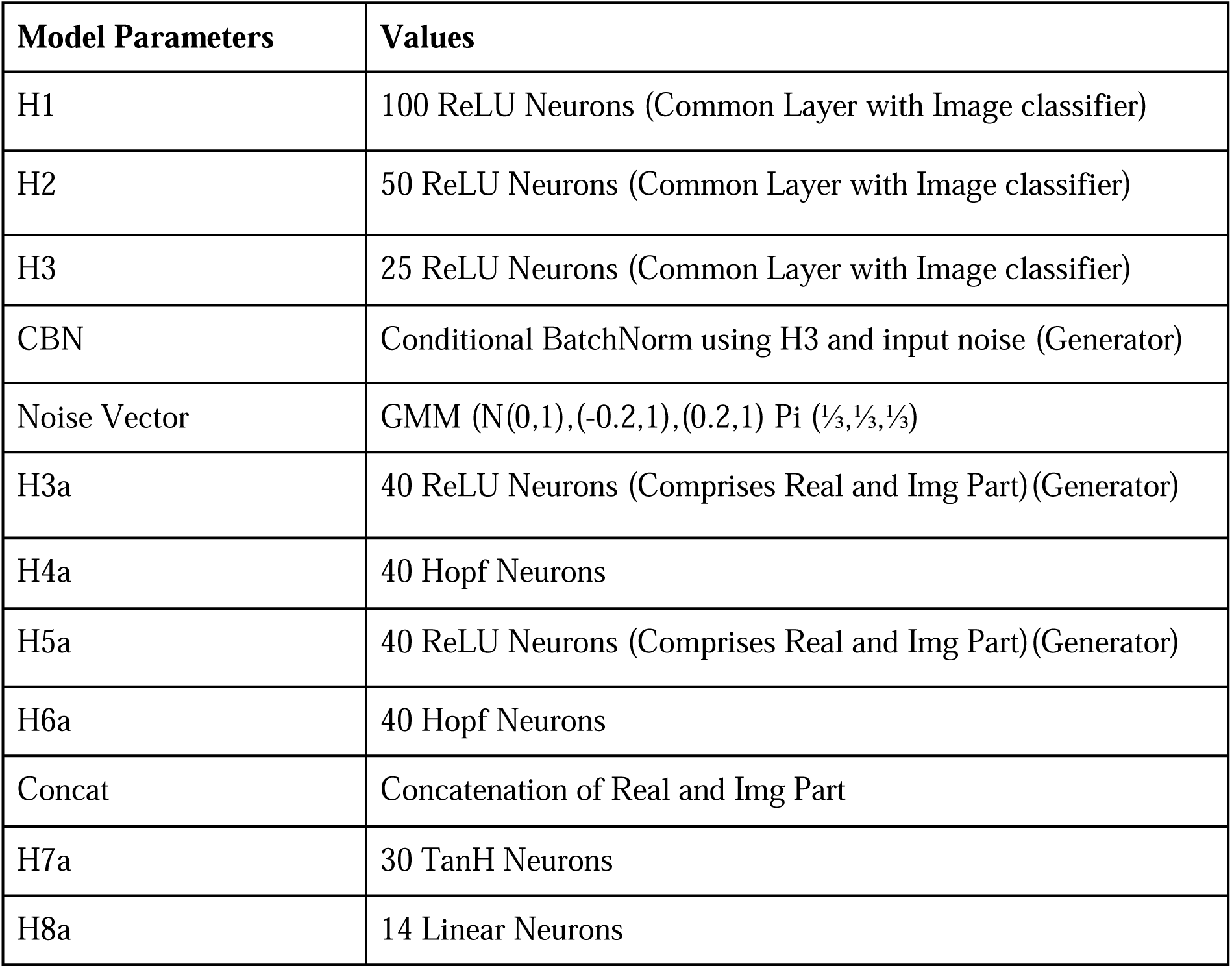
Signal Generator network parameters.

A noise vector, sampled from a mixture of Gaussians, is introduced as input to the next layer after the shared components, following conditional batch normalization using the compressed information from the preceding 25-unit ReLU layer. This generator is responsible for generating EEG signals that correspond to the given visual stimuli.

### 3.3 * Bank of signal evaluator networks

Lastly, the set of K Signal Evaluators takes either the generated or real EEG signals as input. Each classifier consists of two oscillatory layers (comprising a ReLU and Hopf layer), followed by a tanh activation and a final linear layer. The desired output for the signal evaluator network is a ramp signal (Table 6).

**Table 6.** Signal Evaluator network parameters.

| Model Parameters | Values |
| --- | --- |
| C1 | 50 ReLU Neurons (Comprises Real and Img part) |
| C2 | Batch Normalization |
| C3 | 50 Hopf Neurons |
| C4 | 35 ReLU Neurons (Comprises Real and Img part) |
| C5 | 35 Hopf Neurons |
| C6 | 20 TanH Neurons (Comprises Real and Img Part) and Concatenation |
| C7 | Batch Normalization |
| C8 | 10 Linear Neurons |

### 3.4 Network evaluation

#### 3.4.1 Image classification performances

The image classification branch of the classifier-guided network achieved 100% training accuracy. To further test the model, a test font character from Table 2 is passed through the classifier-guided network, which produced both an EEG signal and an image classification output. On the image classification side, the model achieved a 90.9% test accuracy (Table 7). Since the desired output of the image classifier network is a ramp function, the predicted output also follows the ramp target; one example of predicted ramp output is shown in fig 7.

**Figure 7:**
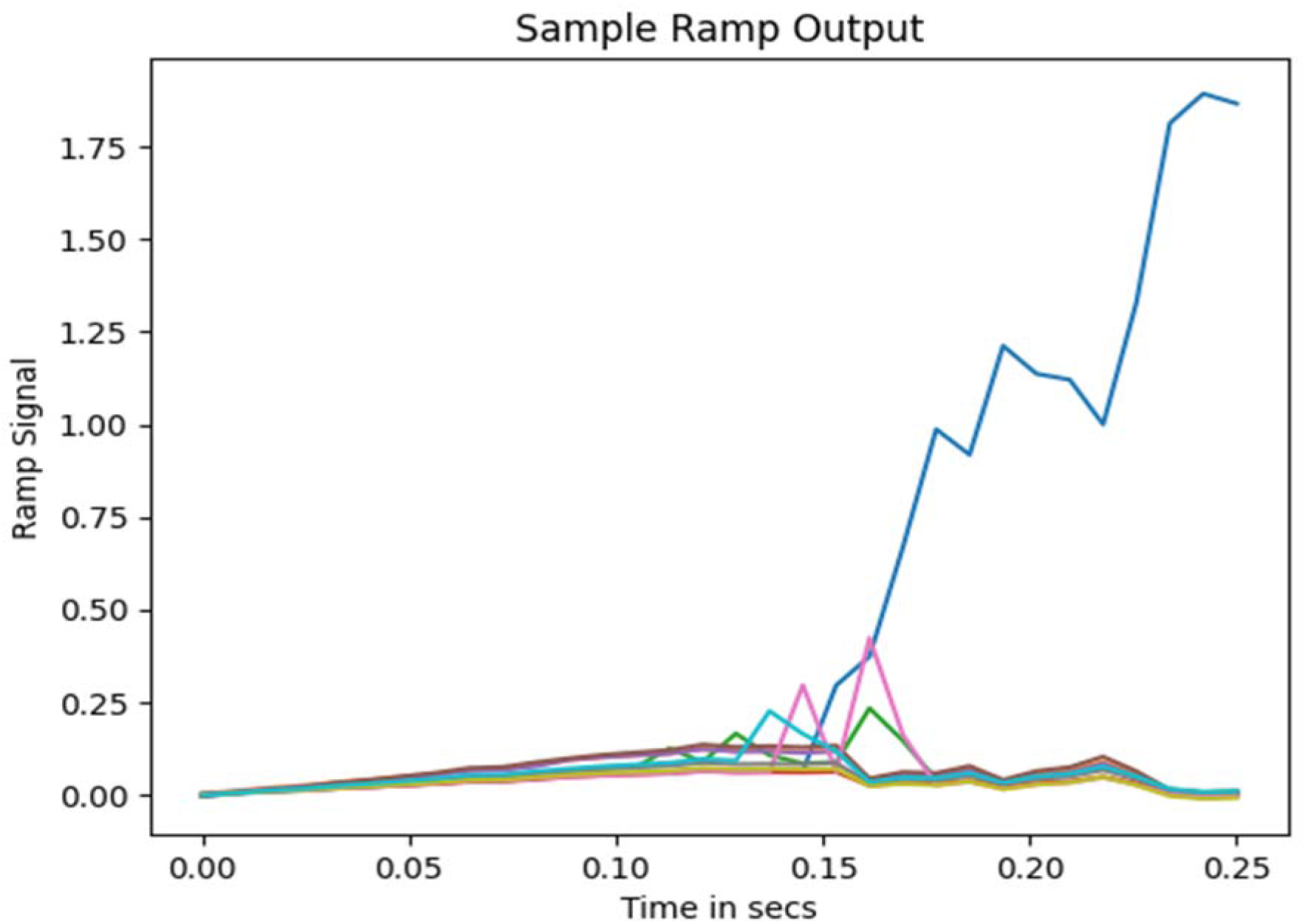
Output predicted during the time when the test image is shown to the model.

**Table 7.** Image classification accuracy using a classifier-guided DONN network.

| Previously Published Work on Imagenet dataset | Accuracy matrices and trainable parameters |
| --- | --- |
| [48 ] model soups, a method to enhance model performance by averaging the weights of multiple fine-tuned models instead of selecting a single best model | 90.98, trainable parameter :2440M |
| [49] MambaVision-L3 | 88.1%, 489.1k |
| [50] SwinV2-G | 90.17%, 3000M |
| Our Proposed (on character Dataset, ImageNet) | Training: 100%, Testing: 90.9, number of epoch: 250, trainable parameter: 205,140 |

#### 3.4.2 Signal classification performances

After training, the EEG data generated by the classifier-guided network during training was evaluated using our pre-trained DONN classifier [42]. In that study, we found out for ThoughtViz dataset DONN provides more accurate result. The classifier achieved 100% training accuracy on a 10-class classification task and 74.05% test accuracy using real EEG data [42]. When the classifier-guided network-generated EEG data (from the training phase) was presented to the pre-trained classifier, it achieved 69.37% classification accuracy (Table 8). Meanwhile, the corresponding generated EEG signal achieved 60.29% classification accuracy when evaluated by the pre-trained DONN classifier [42].

**Table 8.** Evaluation Matrix of EEG signals on a pre-trained classifier.

|  | Accuracy |
| --- | --- |
| Real Signals from Train Data [42] | 100% |
| Real Signals from Test Data [42] | 74.05% |
| Generated Signals from Train Images | 69.37% |
| Generated Signal from Test Images | 60.29% |

#### 3.4.2 Application to Data Augmentation

In this study, we aim to demonstrate the data augmentation capability of our network. For this, we performed few experiments utilising fractions of generated EEG test signals from the Classifier guided DONN model (fig. 2) in retraining the pretrained DONN signal classifier [42]:

a. 90% real EEG data and 10 % generated data
b. 80% real EEG data and 20 % generated data
c. 100% real EEG data and 10 % generated data
d. 100% real EEG data and 20 % generated data

We then tested the accuracy of the retrained DONN module on the test EEG signals we have and observed the classification accuracy rose from ∼75% to ∼83% showing that the generated signals capture remarkable temporal/spectral characteristics of EEG signals corresponding to different classes of images, thus improving the robustness of our signal classifier on unseen test data.

#### 3.4.3 Quantitative measures

Inception score is the one of the popular qualitative measures described in Generative AI literature [65]. It determines the integrity of the data by defining the Kullback–Leibler divergence between the predictions of Inceptionv3 on the generated data and the ground truth labels. In our case, since we have EEG *signal* and Inceptionv3 [90] is a pre trained *image* classifier model, we used our pretrained DONN classifier model [42].

The Fréchet Inception Distance (FID) measures the similarity between two sets of real and synthetic signals [65]. To compute the FID, both the real and generated images are processed through the DONN network [42], and features are extracted from its final layer. Gaussian distributions are then fitted to the extracted features of both real and generated data. The FID score is calculated as the Fréchet distance between these two Gaussian distributions. A lower FID value indicates better quality and greater diversity of the generated images, suggesting they closely resemble the real signals.

#### 3.4.4 Power spectrum analysis

We plotted the generated time series and real EEG time series (fig 8), for different characters. But to judge the quality of the generated data, we compared spectral domain properties of generated EEG and real EEG signal. In order to capture physiologically significant spectral features, we calculated the power within each frequency bands (e.g., delta, theta, alpha, beta, and gamma) (fig. 9(a-e)).

**Figure 8:**
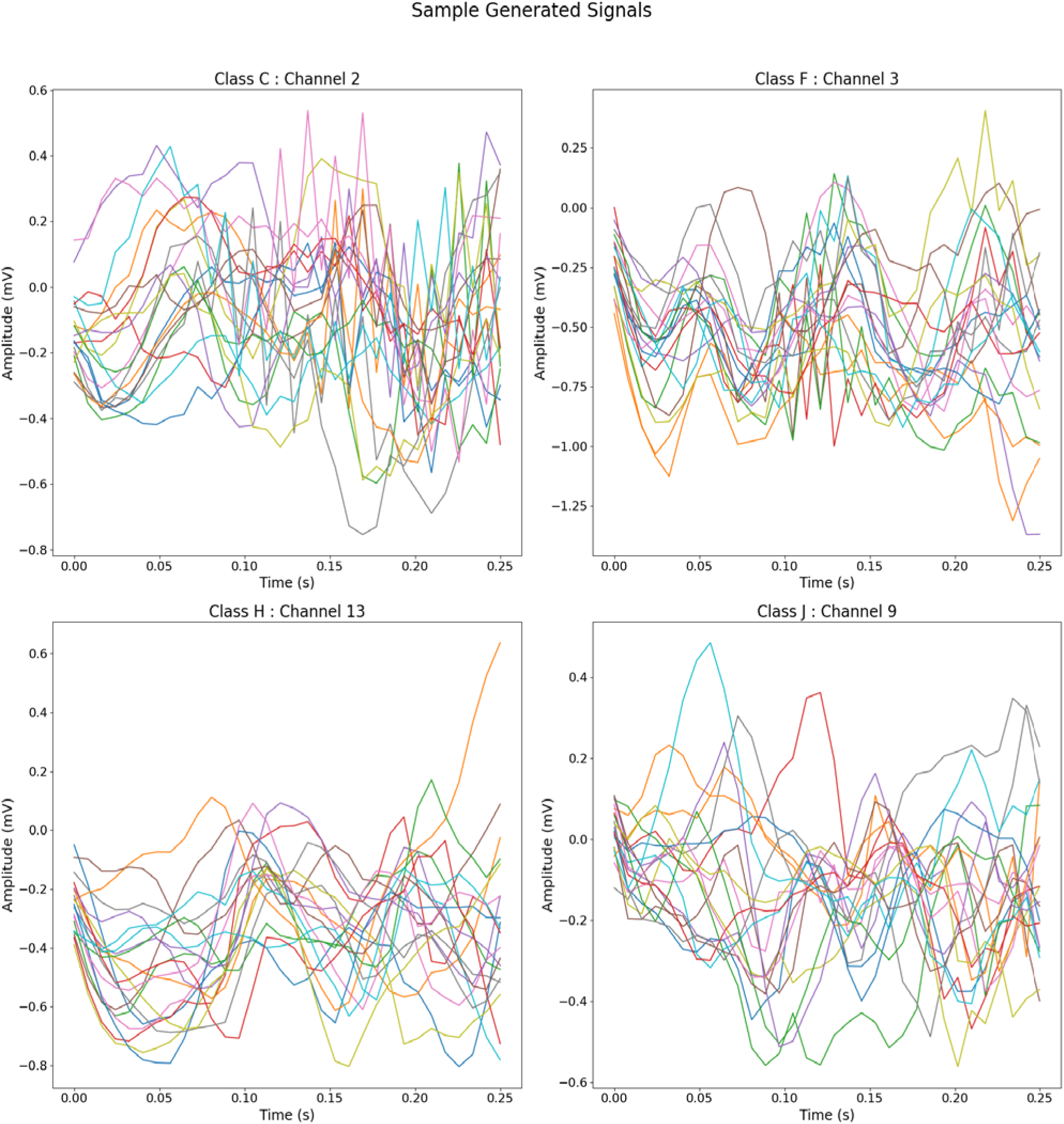
Sample generated signals for different characters

**Figure 9(a-e):**
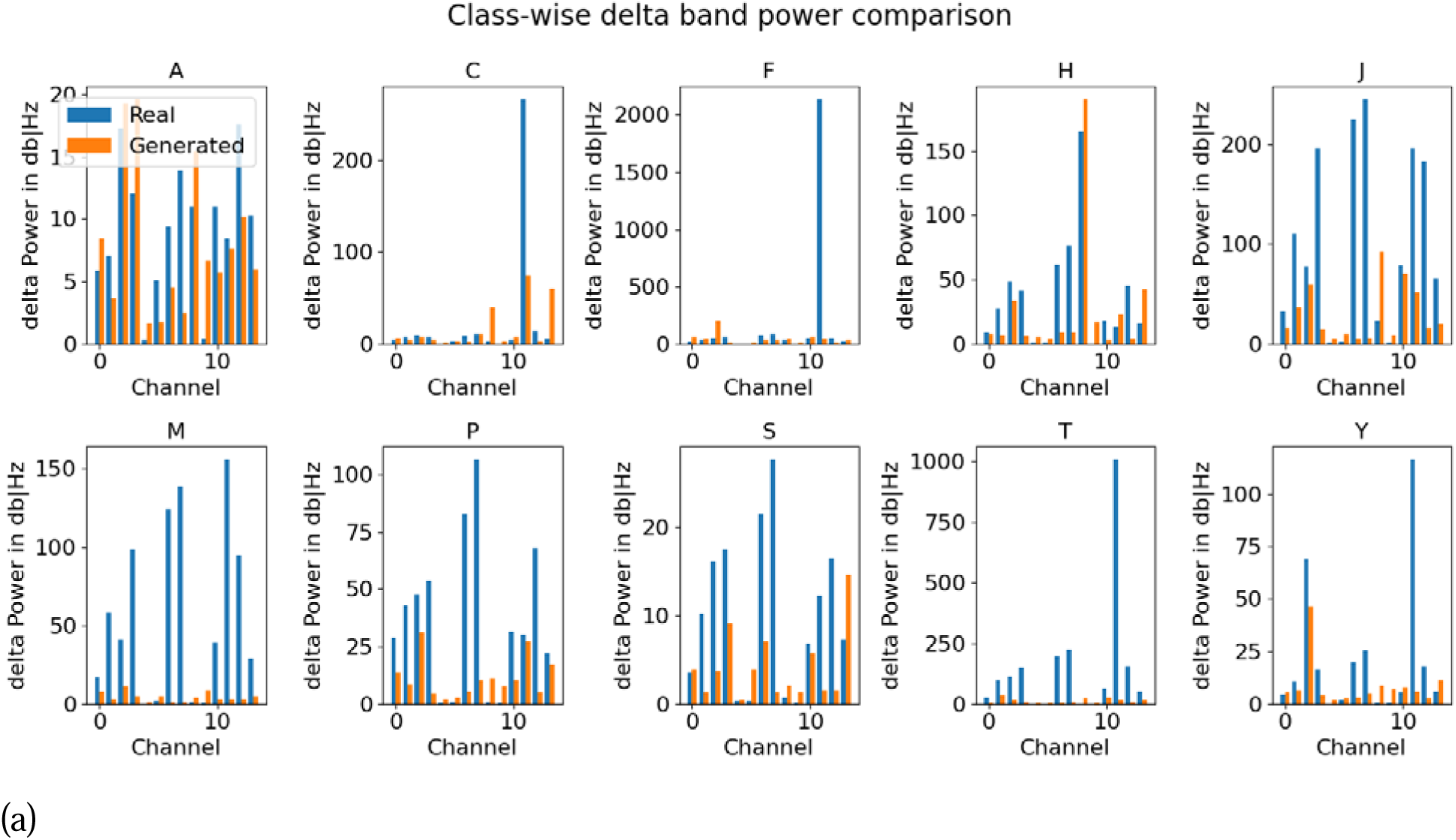
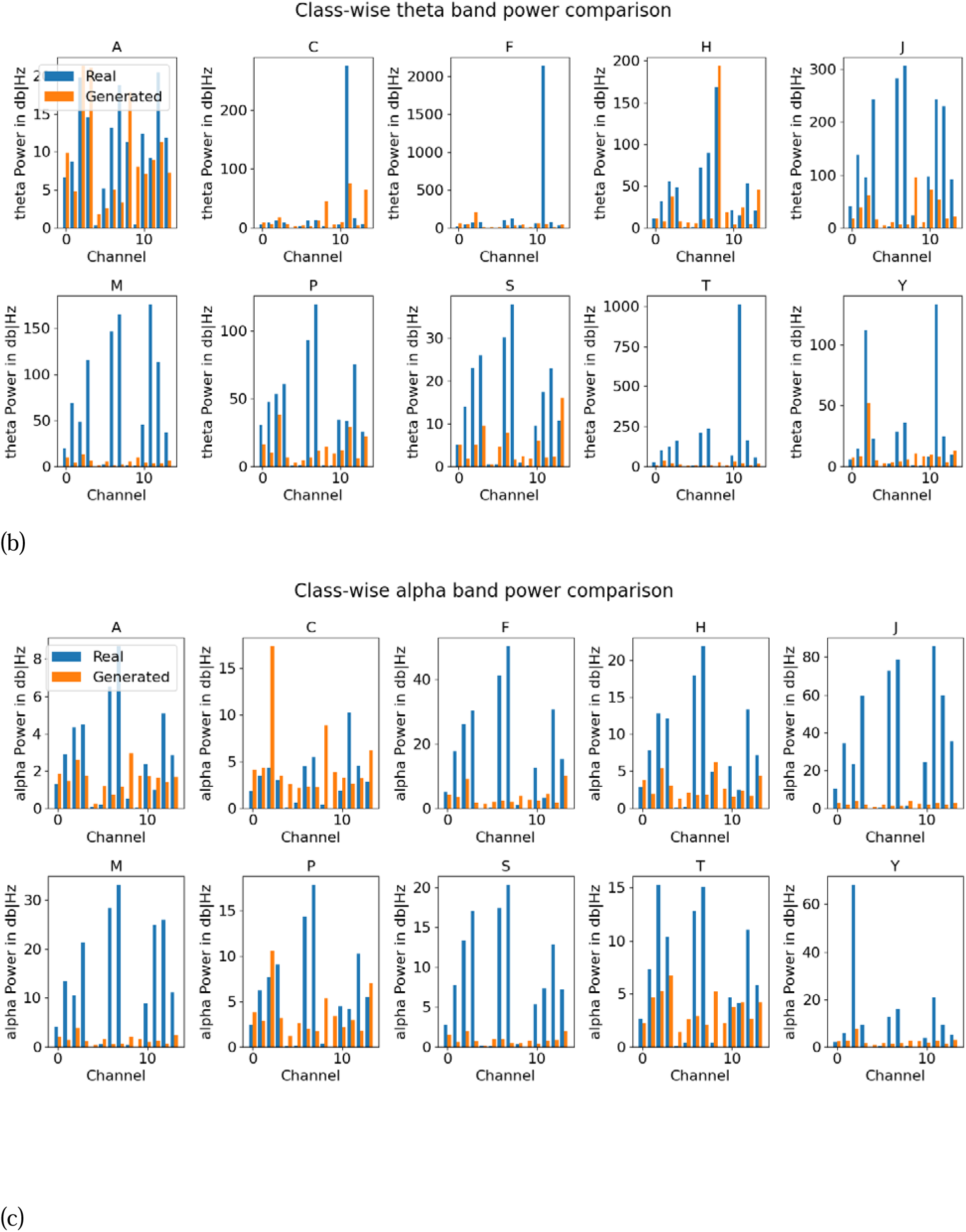
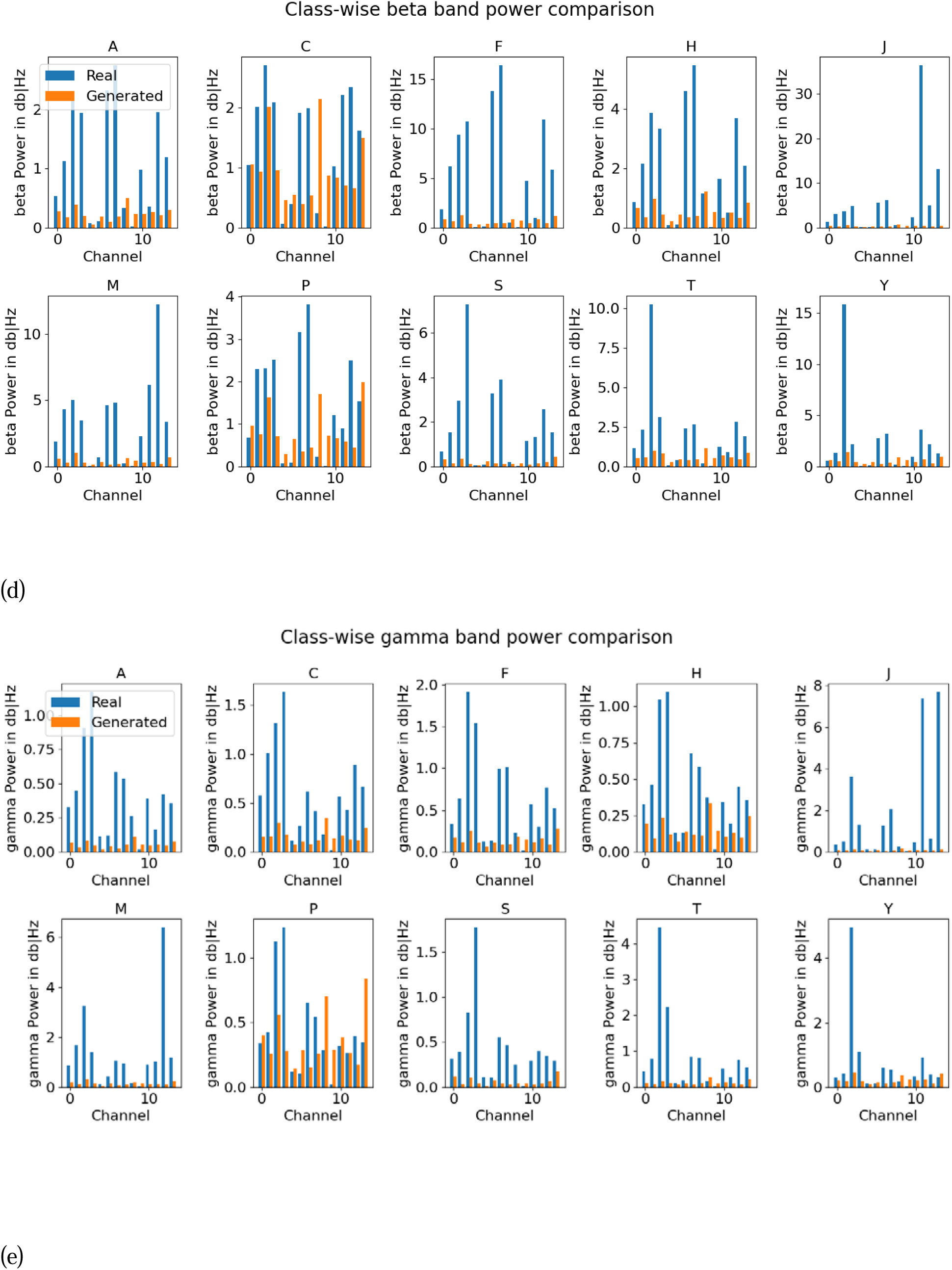
Class*-wise comparison of band power between generated signals (from test image inputs) and real test EEG signals. (a): Delta band, (b): Theta Band, (c): Alpha band, (d): Beta Band, (d): Gamma Band*.

In Fig. (9(a-e)), we present bar plots comparing the power spectra of model-generated signals and real EEG signals across all channels for each character. It can be seen that the generated signals capture the power in the lower-frequency delta band quite well for classes A, P, S, H, and J. Similarly, in the theta band (Fig. 9(b)), the model shows good alignment with real EEG data for classes A, H, and P. Notably, the alpha power in the generated signals demonstrates even stronger correspondence with the real EEG signals—particularly for characters A, C, F, H, P, and T (fig. 9c). The beta band shows moderate agreement for characters C and P, while the gamma band reflects a fair match for characters P and Y in terms of power spectrum features (fig. 9d). Overall, the model captures lower-frequency band power more effectively than higher-frequency components.

We compare the distribution of power spectrum for both real and model-generated signals and the corresponding mean average error is shown in table 11. We show the distribution of power spectrum for channel F4 for class ‘C’ and for channel AF4 for class ‘F’ in figure 10(a-b).

**Figure 10(a-b):**
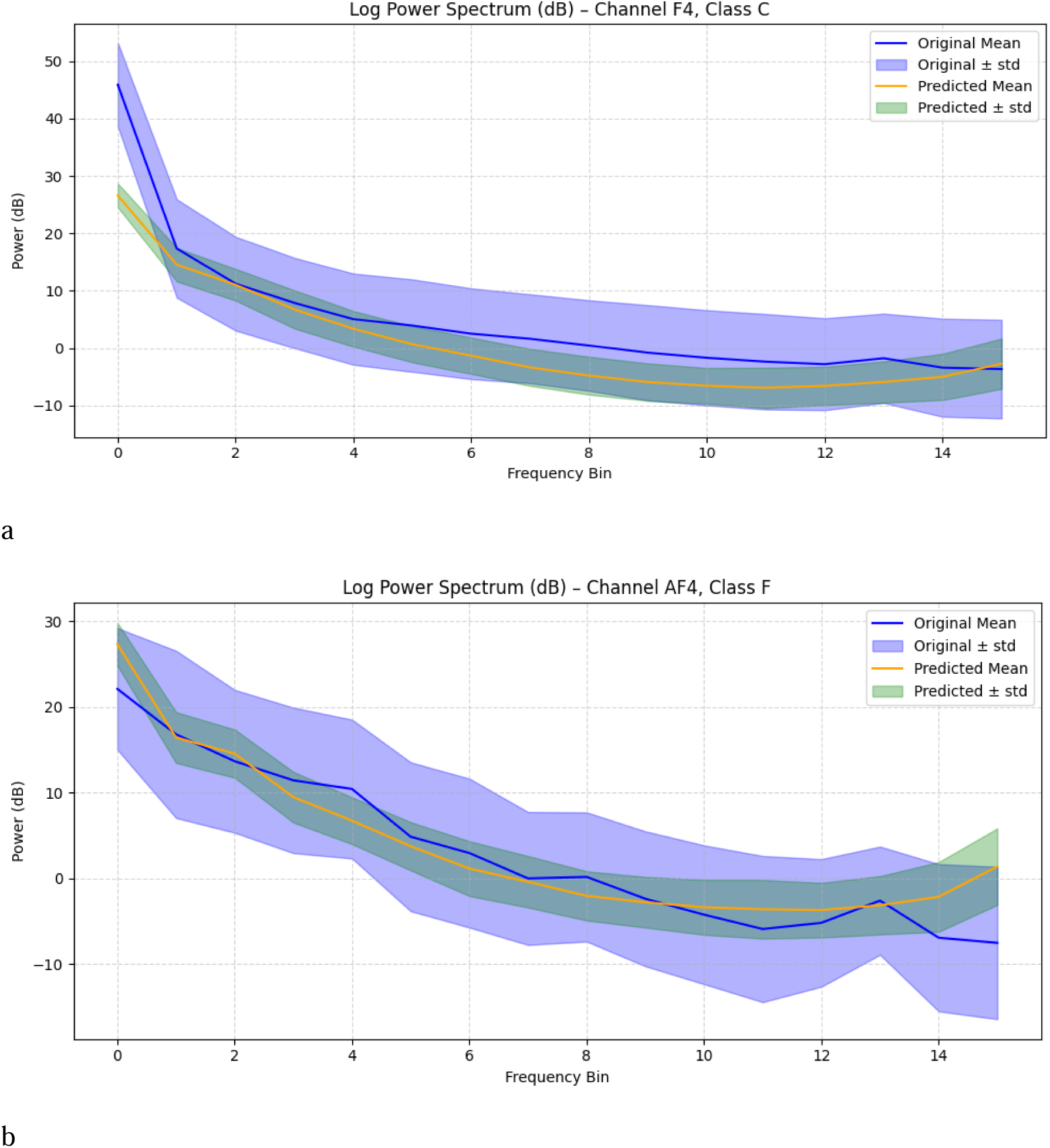
Distribution of model-generated signal power spectrum and desired signal.

**Table 9.** Summary report of Data Augmentation.

| Literature | Accuracy |
| --- | --- |
| [51] BCI competition IV, Dataset 2a, 4 class classification, real data+ 200 trial generated data, model : PP-GAN | $71.03 \pm 3.3$ with SVM |
| [52] BCI 2A Data,2B Data DCGAN-CNN, (Real data+ generated Data) | 2a- increased by 8.7%,2b increased by 12.6 |
| [53] Physionet Sleep Data, RGAN, augmented with 50% real data | +39.1%, improved than original ($93.4 \pm 4.2$) |
| Our proposed model: Thought Viz<br>10-class classification, Hybrid DONN | 90% real EEG Data+ 10% generated data-82.85<br>80% real EEG Data+ 20% generated data-83.93<br>100% real EEG Data+10% Generated EEG-82.68<br>100% real EEG Data+ 20% Generated EEG-82.70 |

**Table 10.** Comparison of IS, FID metrics across other works.

| Works | Model | Dataset | Class | Result |
| --- | --- | --- | --- | --- |
| [54] | WGAN<br>+<br>(TSF-MSE)<br>loss<br>function | Action<br>observation<br>dataset | 4-class | 67.67 |
|  |  | GAL dataset | 6-class | 73.89 |
|  |  | Motor<br>Imagery<br>(BCI) | 4-class | 64.01 |
| [55] | **cVAE- GAN** | BCI-2A<br>Dataset | **4-class** | IS-1.35<br>FID-11.36<br>SWD=0.067 |
| [56] | WGANs |  |  | IS=1.36<br>FID: 9.523 |
| [57] | Sig-GAN | Publicly<br>Sleep-EDF<br>database<br>(PSG) test |  | IS=0.51<br>FID=39.53 |
| Our proposed<br>model | Classifier<br>guided<br>DONN | ThoutViz<br>Visual EEG<br>Data | 10 Class | IS-1.45<br>$\pm 0.16$<br>FID: 7.21 |

**Table 11.** Class wise Power spectrum Mean Average Error (MAE) between generated and real EEG signals.

| Class | MAE across channels |
| --- | --- |
| Class A | 5.48 |
| Class C | 5.97 |
| Class F | 6.69 |
| Class H | 5.99 |
| Class J | 7.90 |
| Class M | 7.18 |
| Class P | 7.15 |
| Class S | 7.29 |
| Class T | 5.81 |
| Class Y | 6.72 |

We conducted a comparative analysis of the topographic power distributions across frequency bands for Class C, using both real EEG data (fig. 11(a) and model-generated EEG data (fig. 11(b)).

**Figure 11(a-b):**
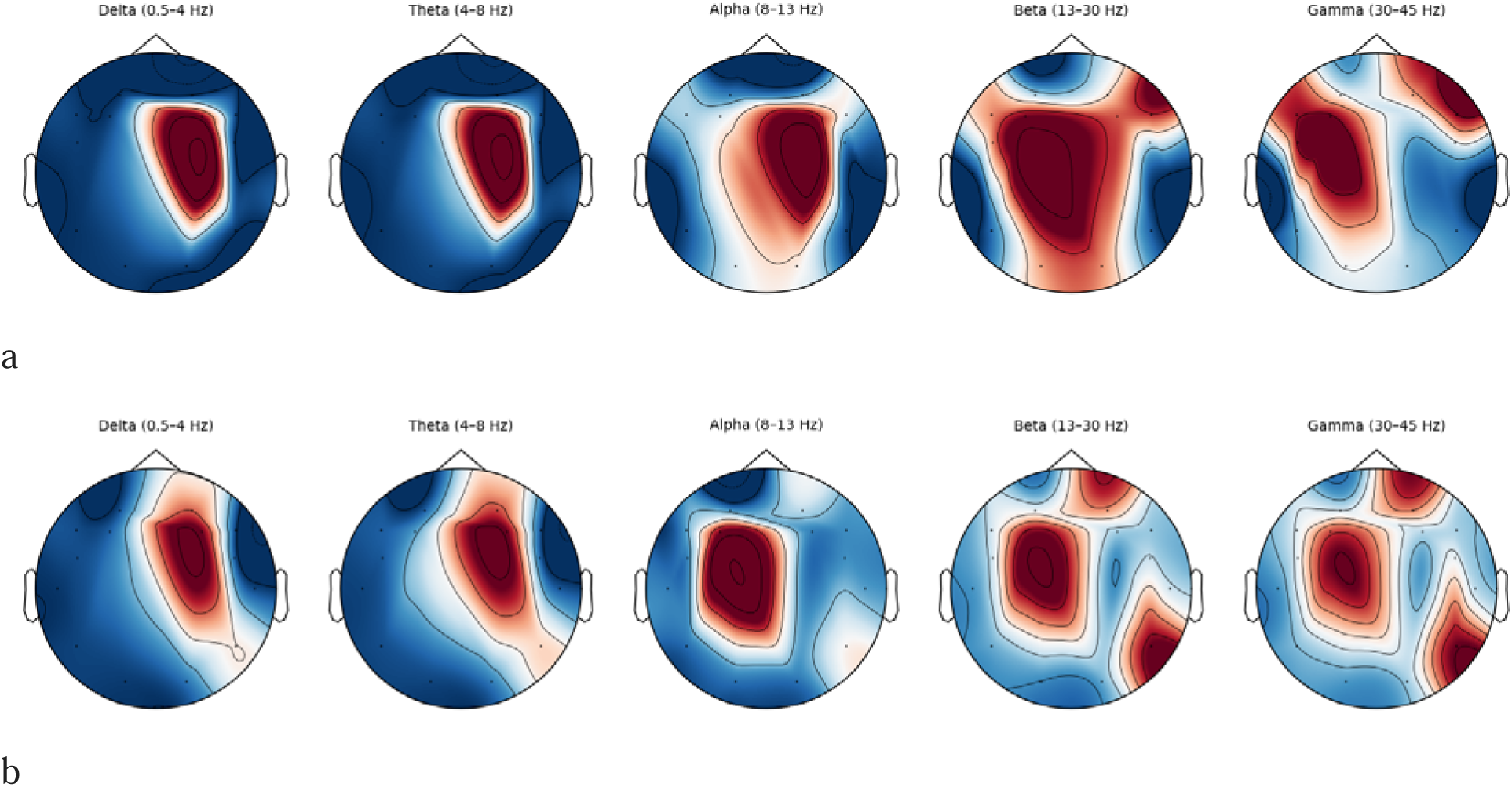
Topographic Representation of Class C Band Power on (a): Real EEG,(b): Generated Data.

In the Delta band, both real and generated EEG signals exhibit strong power concentrated in the left-central region, extending toward the midline, indicating that the model effectively captures the characteristic low-frequency spatial pattern of Class C.

In the Theta band, real EEG data shows a power peak slightly anterior to the delta hotspot, still within the left-central area. The generated EEG presents a similar but more diffuse distribution, with power oriented in the same region, suggesting a reasonable approximation of theta dynamics.

For the Alpha band, both real and generated signals show a strong concentration of power over the central and midline parietal regions, demonstrating a high level of spatial agreement.

In the Beta band, the real EEG shows power localized in the central and parietal areas, whereas the generated EEG displays a more widespread distribution, though it still reflects some bilateral symmetry seen in the real data.

In the Gamma band, the alignment between real and generated data is less pronounced, with no clear spatial matching.

Overall, the model-generated EEG for Class C demonstrates good spatial correspondence with the real EEG in the Delta, Theta, and Alpha bands, with the Beta band moderately well captured. The Gamma band, however, shows limited agreement.

## 4. Discussion

In this study, we introduce a Classifier guided Deep Oscillatory Neural Network (cDONN) capable of both multimodal signal generation and image classification. The image classification performance of our model is benchmarked against existing literature, as shown in (Table 7), demonstrating its accuracy advantage. Additionally, we provide a comparative analysis of signal generation quality using Inception Score and FID metrics (Table 10), where our model exhibits notable improvements over previously reported methods.

In our earlier work, we developed a purely generative model based on an oscillatory network comprising Hopf oscillators, capable of predicting and generating EEG time series across different sleep stages [10]. Building on that foundation, we extended the Hopf-based generative framework into DONN, an oscillatory input-output architecture, capable of mapping input time-series onto output time-series [44]. This is an important result in computational neuroscience where is currently a race to build models that can simultaneously learn behavior and model brain dynamics.

To the best of our knowledge, this work represents the first attempt to develop a model that simultaneously captures both behavioral and dynamical aspects of brain-inspired computation within the context of image classification. This biologically motivated approach marks a significant shift from traditional deep learning methods, which primarily focus on maximizing classification accuracy without considering the underlying neural dynamics.

Image classification is a fundamental and widely explored application area in the field of machine learning. Over the years, several high-performing deep learning architectures such as AlexNet [58], GoogLeNet [59], VGG16 [58], and DenseNet [60] have been extensively adopted for tackling image classification tasks. On the Char74K dataset, the VGG-16 model demonstrated a high classification accuracy of 95.62% for recognizing 26 alphabetic characters and 10 numeric digits [61], while another study introduced a quantum-inspired CNN to classify 10 classes with 90% accuracy [62]. Additionally, the ResNet architecture was employed to classify 26 lowercase alphabet characters, achieving an accuracy of 81.81% [63]. A. Botalb et al. [64] further reported accuracies of 89.47% and 92% using MLP and CNN models, respectively, on the EMNIST dataset for 26-class handwritten character recognition. Moreover, a novel unsupervised feature selection (UFS) combined with a Multi-Support Vector Machine (MSVM) framework has been proposed to enhance handwritten character recognition across multilingual scripts, including Kannada, Arabic, and English [65]. These models have consistently demonstrated outstanding accuracy and robustness, making them the go-to solutions for many real-world applications.

While these architectures excel at behavioral performance, our research takes a different direction by aiming to design a model that not only performs well in classification tasks but also emulates the dynamics of the biological brain. The primary objective of our model is to integrate both behavioral output and dynamic properties, thereby creating a more biologically inspired framework that aligns closer to how the human brain processes visual information.

Although our classifier may not outperform some of the state-of-the-art models – which only perform image classification but do not seek to model brain dynamics – like those mentioned above in terms of raw accuracy (Table 7), it still delivers competitive and meaningful results, especially when considering the added value of its neural dynamics and interpretability.

Electroencephalography (EEG) offers a high-temporal-resolution window into brain activity but faces challenges like low signal-to-noise ratio, non-stationarity, and inter-subject variability due to anatomical and procedural differences. These issues hinder the generalizability of deep learning models, which also struggle with limited labeled data common in neuroscience. Data augmentation has emerged as a promising solution, expanding training sets with synthetic data to improve model robustness. While well-established in vision and NLP, EEG-specific augmentation remains underexplored. Some notable efforts include: Dai et al. [49] using hybrid-scale CNNs with time and frequency domain recombination, yielding 2–3% accuracy gains; VAEs achieving ∼3% improvement [66]; and GAN-based methods, including DCGANs on EEG spectral images, enhancing classification by 3.22–5.45% [67]. Our cDONN model surpasses these approaches (Table 9), highlighting its strong potential in EEG augmentation.

Furthermore, in Table (7), we present a comparison of the number of trainable parameters between our proposed model and the standard architectures. This comparison highlights that our model is significantly more lightweight in terms of complexity, yet manages to deliver promising performance. This balance between computational efficiency and biological plausibility positions our model as a strong candidate for future work in neuromorphic computing and brain-inspired artificial intelligence.

Looking ahead, a key direction for our future research is to integrate more complex and powerful architectures—such as AlexNet, GoogLeNet, VGG16, and other deep learning models—into the classification component of our system. These architectures have demonstrated strong performance on standard image classification benchmarks and, when appropriately incorporated, could further enhance the behavioral accuracy of our model. However, it is important to highlight a major distinction between our work and previous studies. Most existing deep learning models are trained on large-scale image datasets such as ImageNet [41], MNIST [68], and Char74K [69], which contain hundreds of thousands to millions of labeled samples. These datasets are purely visual in nature, and models trained on them do not need to account for additional modalities like brain signals.

In contrast, our approach is fundamentally different: we are working with multi-modal data, which includes both images and the corresponding EEG signals generated when a subject views those images. Specifically, we use the ThoughtViz dataset [40], where the authors introduced 800 font variations for a single character class. For our training purposes, we are limited to only 800 training images, each associated with its own EEG signal recording.

This relatively small dataset size presents a significant challenge for traditional deep learning models, which typically require large volumes of data to achieve high classification accuracy. As a result, while our model may not currently match the performance of state-of-the-art image classifiers on standard datasets, it still delivers notable results given the limited data and added complexity of EEG integration.

In contrast, our approach is fundamentally different: we work with multi-modal data that includes both images and the corresponding EEG signals recorded as subjects view those images. Specifically, we utilize the ThoughtViz dataset [40], in which the authors introduced 800 font variations for a single character class. For our training, we are limited to just 800 images, each paired with its corresponding EEG recording. This relatively small dataset poses a significant challenge for traditional deep learning models, which typically require large amounts of data to achieve high classification accuracy. Consequently, while our model may not yet match the performance of state-of-the-art image classifiers on standard datasets, it achieves promising results given the limited data and the added complexity of EEG integration.

In addition, integrating a reinforcement learning framework into the cDONN model could significantly enhance its biological plausibility by enabling adaptive learning mechanisms akin to those observed in the brain.

## 5. Conclusion and future direction

In this work, we present a novel classifier guided Deep Oscillatory Neural Network (cDONN) framework that uniquely integrates image classification with biologically inspired EEG signal generation, effectively bridging the gap between behavioral interpretation and neural dynamics within a single architecture. By leveraging the rich dynamics of Hopf oscillators, the cDONN model demonstrates the capacity to emulate both cognitive and neurophysiological processes, offering a dual-output system that aligns more closely with how the brain functions. While our classifier may not yet outperform traditional deep learning architectures in raw accuracy, it provides meaningful results under limited data conditions and offers a biologically grounded alternative with significantly reduced computational complexity.

Our results show strong coherence between generated and real EEG signals, particularly in key frequency bands and cortical regions, validating the fidelity and realism of our generative component. Additionally, benchmarking against traditional evaluation metrics such as Inception Score and FID further substantiates the quality of the synthesized EEG data.

This research introduces a paradigm shift in multi-modal learning by combining perception (via image classification) and cognition (via EEG dynamics) into a unified, interpretable model. Looking ahead, we aim to incorporate more powerful classification architectures and expand the dataset to improve performance and generalizability. Our approach opens new directions for future work in neuromorphic computing, neuro-symbolic AI, and human-computer interaction by paving the way for systems that both understand and replicate brain-like behavior.

## Acknowledgement

We acknowledge the financial support from the Parkinson Therapeutics Lab, IIT Madras. The authors acknowledge the support of fellow intern Talakanti Sravan Kumar Reddy for his help in developing of the pre-trained DONN model.

## Author Contribution

**Sayan Ghosh:** Methodology, software, Data curation, Investigation, Formal analysis, Validation, Writing-Original Draft, Writing-Review & Editing, Conceptualization. **Sapna Raja**: Methodology, software, Data curation, Investigation, Formal analysis, Validation, Writing-Original Draft, Writing-Review & Editing, Conceptualization. **V. Srinivasa Chakravarthy**: Project administration, supervision, Funding acquisition, Writing-Review & Editing, Conceptualization.

